# Developmental Connectomics of the Mouse Cerebellum

**DOI:** 10.1101/2025.09.15.676403

**Authors:** Nagaraju Dhanyasi, Yaron Meirovitch, Vikrant Kapoor, Alyssa M. Wilson, Shuohong Wang, Donglai Wei, Richard Schalek, Rong Ma, Yuelong Wu, Akhila Manthena, Diego Cordero Cervantes, Zudi Lin, Yunlei Liu, Sanjana Chimaladinne, Benjamin Morris, Indu A. Vaddiparti, Yan Zhu, Sanjana Dokiburra, Riti Parampalli, Fuming Yang, Daniel R. Berger, Hanspeter Pfister, Venkatesh N. Murthy, Jeff W. Lichtman

## Abstract

To uncover the developmental processes that establish the precise patterns of synaptic connectivity in the CNS, we employed a connectomic approach in the mouse cerebellar cortex between birth and 2 weeks of age. There were dramatic quantitative and qualitative changes in the structure and connectivity of cerebellar cells. Parallel fiber synapses onto Purkinje cells increased ∼500-fold, with the most rapid growth taking place a week after birth. To support this profound synaptogenesis, Purkinje cells generated thousands of transient parallel fiber-oriented filopodia that received nascent synapses from parallel fibers. Importantly, we find that granule cells initiate synaptic output onto Purkinje cells only after receiving mossy fiber input, revealing a sequential, input-dependent logic for circuit assembly. In sharp contrast to the concurrent pruning of climbing fiber inputs, parallel fiber connectivity expanded and became highly individualized during development. Despite anatomical overlap, neighboring Purkinje cells share significantly fewer parallel fiber inputs than expected by chance. Moreover, parallel fibers themselves diverged spatially, further enforcing selective input allocation and resulting in highly specific parallel fiber cohorts for each Purkinje cell. Our findings uncover a mechanistic sequence in which early afferent activity and transient cellular structures guide the selective wiring and expansion of parallel fiber input to Purkinje cells, establishing developmental principles that ensure functional specificity in the mature brain.

## Introduction

Each neuron in the brain receives and transmits signals to thousands of other neurons via synapses, forming a dense network of neurons that underlies brain function ^1–4^. However, the precise details of the connectivity are poorly understood. There are a number of challenges to comprehending the organization of these connections. First, in most parts of the brain a significant proportion of the input to and output of a neuron is distributed over large areas, making it difficult to identify both the origins of the recipient synapses and the synaptic targets of a neuron. Second, the diversity of neuronal cell types in most brain regions is large and not fully defined. Thus, finding circuit principles is confounded by the variety of circuits in a brain region ^5–11^. Third, the pattern of connections is probably not stable. Synaptic plasticity in adults and certainly major connectivity refinements in development are a challenge to successfully describing synaptic circuits. This is especially relevant to mammalian nervous systems where learning and experience are known to modify connectivity ^12–23^. Therefore, mapping developmental trajectories of neuronal networks is a critical aspect for reaching an understanding of how mature circuits come into existence ^24–26^. Longitudinal studies of brain connectivity may also be valuable to understanding the origins of miswiring in neurodevelopmental disorders ^27–31^

To mitigate the challenges of the study of circuits mentioned above, we chose to investigate the development of connectivity in the cerebellum, known for its highly stereotyped modular organization ^32–36^. The cerebellum contains a canonical circuit structure that repeats millions of times throughout its extent. This iterative structure allows for comparison of circuits across development. In addition, some of the cells, most notably the granule cells and some of the inhibitory cell types, are completely contained within the cerebellar cortex so that all their input and output connections are available for study. Finally, because of the modular structure, many connected cells can be analyzed in relatively small volumes. In this work we have focused our attention mainly on the synaptic connectivity of parallel fibers (the axons of granule cells) and their postsynaptic targets, Purkinje cell dendrites. Purkinje cells provide the only output axons that exit the cerebellar cortex. The granule cells are the most numerous cells in the mammalian brain ^34^ and they make a remarkably large number of synapses on Purkinje cells. Because mature Purkinje cells possess a large, fan-shaped dendritic arbor ^37–39^ that receives tens of thousands of synapses from parallel fibers, owing to its topography, this circuit allows for uncovering the principles of the organization of patterns of connections in neural circuits ^35,40,41^. Millions of parallel fibers run orthogonally to the 2D dendritic trees of Purkinje cells resulting in what appears to be a crystalline organization ^42–46^. The dense packing of parallel fibers, which are individually smaller than the diffraction limit of light microscopy, poses a significant challenge for revealing their connection patterns using conventional light microscopy techniques. We thus used serial section electron microscopy which allowed straightforward inquiry into questions like: which parallel fibers are connected to which Purkinje cells and how do the specific patterns emerge during development?

We studied the cerebellum at five different stages of postnatal development by generating volumetric data sets at postnatal day (P) 0, P3, P7, P10 and P14. This longitudinal analysis revealed considerable complexity in the steps that lead to the standard adult circuitry and neuronal structure of this well-known brain region.

## Results

To study the maturation of parallel fiber connectivity to Purkinje cells during the first two weeks of postnatal life, we used serial section electron microscopy to reconstruct Purkinje cells and cohorts of parallel fibers innervating Purkinje cells at P0, P3, P7, P10 and P14 (Figs. S1 A-E) (see Methods for how Purkinje cells were identified at P0-P7). Reconstructed Purkinje cells at each age showed obvious differences in their appearance (Fig. 1A-E, Videos 1-4, and Fig. S2 A-R; See Fig. S3, for confocal images of Purkinje cells at P6 and P10).

**Figure 1.**
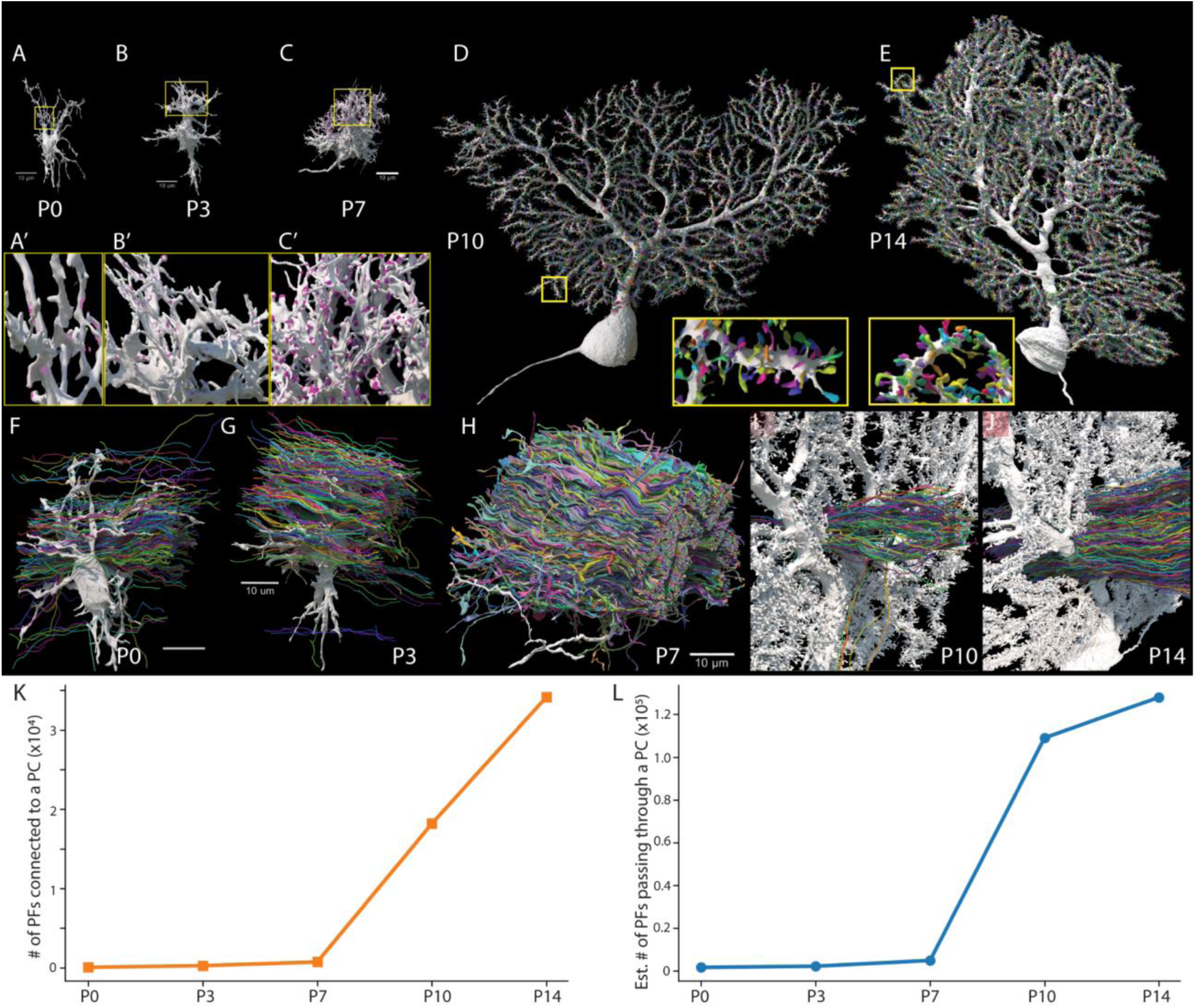
Profound changes in Purkinje cell morphology and connectivity in the first two postnatal weeks. A-E) Reconstructions of Purkinje cells at 5 time-points between birth and 14 days of age, at the same size scale. Insets A’, B’, and C’ show high magnification blow-ups of the yellow boxed regions in panels A, B, and C. Sites of parallel fiber synapses onto the Purkinje cells are shown as pink patches. In panels D and E, the Purkinje cell spines are tinted in random colors. F-J) Shown are clusters of adjacent parallel fibers passing through the dendritic arbors of an individual Purkinje cell at each of 5 ages between P0-P14. K) Graph showing the number of parallel fibers innervating a Purkinje cell, which ranges from 64 at P0 to 34,178 at P14. L) An estimate of the total number of parallel fibers passing through a Purkinje cell dendritic arbor, ranging from 1,645 at P0 to 124,615 at P14. Scale bars: 10 µm.

### A 500-fold increase in parallel fiber synaptic input to Purkinje cell during the first two postnatal weeks

To explore how the characteristic changes in Purkinje cell dendritic morphology over the first two weeks of postnatal life are related to the maturation of parallel fiber innervation onto Purkinje cells, we quantified parallel fiber input to individual Purkinje cells at P0, P3, and P7. Owing to the immense number of innervating parallel fibers during the second postnatal week, we then limited our reconstructions to several hundred axons (343) at P10 and several thousand at P14 (∼3000) (Fig. 1 F-J, videos 5-8). In addition, we reconstructed all the spines on one P10 and one P14 Purkinje cell to get a better estimate of the exact number of parallel fibers innervating a Purkinje cell (Fig. 1 D and E).

This longitudinal analysis showed a two-phase synaptogenesis process for parallel fibers. During the first postnatal week, the number of parallel fiber axons that made histologically unambiguous synapses with a Purkinje cell increased from 64 at P0 to 271 at P3, and to 742 at P7. This amounts to a 10.5-fold increase in the number of parallel fiber axons converging on a Purkinje cell during the first postnatal week. After P7 this convergence rate accelerated further. At P10 we reconstructed all the spines of a Purkinje cell and found 18,700 spines in total. We expected that the majority of these spines would be innervated by parallel fibers. To check this, we identified the inputs to each of these spines by synaptic structure. 340 of the spines (1.8%) were innervated exclusively by climbing fibers. The rest were innervated by a combination of excitatory (parallel fiber) and inhibitory (basket and stellate cells). To detail the inhibitory input more completely, we examined the source of innervation to 1000 spines in the proximal dendritic tree. We chose these because they are the most mature spines on the Purkinje cell ^47,48^. We found 9 inhibitory axons. Two of these shared a spine with a parallel fiber, and the other 7 were the sole input to a spine (data not shown). These 9 inhibitory axons were mainly branches of basket cell axons that also innervate the somata of Purkinje cells. These results suggest that approximately 97% of the spines are innervated by parallel fibers (18,195 of 18,700 spines). This large number of parallel fibers equates to a 24.2-fold increase from P7 to P10 (Fig.1K).

From P10 to P14, parallel fiber connectivity continued to increase. We counted 35,748 spines on a P14 Purkinje cell; of these, 2.1% (750 spines) were innervated by climbing fibers, 2.3% 820 spines) by local inhibitory interneurons and a total of 34,178 (95.6%) of the spines were innervated by parallel fibers, a further ∼1.9-fold increase (Fig. 1K). In summary, while there was a 10-fold increase in parallel fiber axon convergence during the first postnatal week, there was a massive 46.9-fold increase during the second postnatal week. This leads to a 533-fold increase in the number of innervating parallel fibers per Purkinje cell over the first two postnatal weeks.

### An accretive model of formation for the assembly of parallel fiber to Purkinje cell connectivity

We identified several processes responsible for the large increase in the number of parallel fiber axons innervating Purkinje cells in the second postnatal week. First, there was a substantial increase in the number of parallel fibers passing through Purkinje cells between P7 and P10 (3,780 to 140,000, a ∼47-fold increase; Fig. 1L). This dramatic addition of parallel fibers is likely related to the wave of granule cell neurogenesis in rodent early postnatal life ^49–51^, in which new granule cells quickly establish parallel fibers. Second, these new parallel fibers, which are located in the more superficial parts of the molecular layer (near the pia), were joined by the upward-growing dendrites of Purkinje cells that invade their territory (Fig. S2S; consistent with previous reports ^38,52,53^ and fan out by adding side branches that overlap with progressively larger numbers of parallel fibers (Fig. 1D and E and Fig. S2 O-R)). We found these newer sites of contact, at least at the outset, to have a lower density of parallel-fiber-to-Purkinje-cell-dendrite synapses (see Fig. 2 H, I and Fig. S4H). Third, there was an increase with age in the probability that a parallel fiber passing through a Purkinje cell dendritic arbor would establish a synapse. At P0, only 4.3% of the parallel fibers passing through a Purkinje cell formed synapses at P3 12% of parallel fibers established synapses on Purkinje cell dendrites in their immediate vicinity, 21% did at P7, 20% established synapses at P10, and 25% at P14 (see Fig. S4E). Notably, at all five developmental stages parallel fibers mostly (90%) established one synapse per Purkinje cell they innervated (Fig. S2T). This single-synapse motif is similar to what has been observed in the adult cerebellum ^41–44^ and indicates that parallel fiber synapse addition entails an increase in “fan-in” (the number of unique afferent parallel fibers innervating each Purkinje cell).

**Figure 2.**
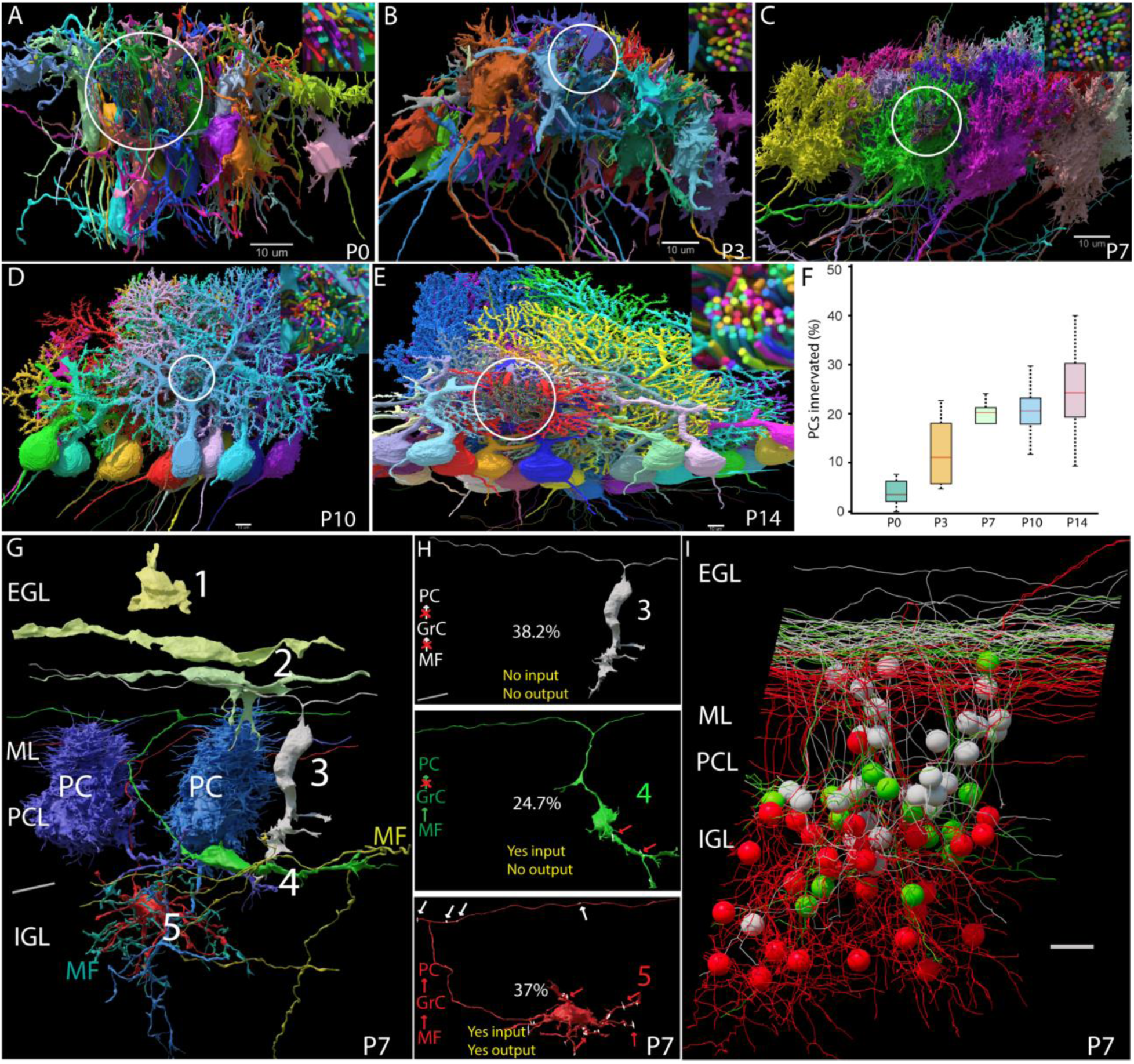
Developmental changes in the structural organization and connectivity of sets of adjacent Purkinje cells. A-E) Shown are the progressively more orderly arrangements of Purkinje cell dendrites and the location of their soma over the first two postnatal weeks. Cohorts of parallel fibers (within the yellow circles) that innervate one or more of the multicolored Purkinje cells are reconstructed at each age. F) Boxplot showing the increasing fraction of Purkinje cells innervated by each parallel fiber between P0 and P14. G-K) Sequential assembly of granule cell input and output synaptic connectivity. G) Granule cells at five maturational stages (labeled 0-4) from incipient (1) to bipolar (2), to migratory (3), to later migratory (4) to mature (5). H) Panels showing three connectivity stages for individual granule cells and the percentage breakdown of the three connectivity classes of granule cells: Top panel shows a migrating granule cell (such as cell 3 in G) whose dendrites did not receive synapses from mossy fibers and whose parallel fiber did not establish synapses onto Purkinje cells. The middle panel shows a granule cell that is completing its migration (such as cell 4 in G). Its dendrites received synapses from one or more fibers (red arrows), but its parallel fiber did not yet establish any outgoing synapses. The bottom panel shows a granule cell that has completed its migration. This cell is both innervated by mossy fibers (red arrows), and its parallel fibers have established several connections with Purkinje cells (yellow arrows). I) A reconstruction showing the different locations of the three connectivity classes of granule cells (white, green, and red cells). EGL: External granule cell layer, ML: Molecular layer, PCL: Purkinje cell layer, IGL: Internal granule cell layer, PC:Purkinje cell, GrC: granule cell, MF: Mossy fiber. Scale bars: 10 µm.

To further assess how this progressive accretion of parallel fiber-to-Purkinje cell synapses impacted local circuitry, we also assessed developmental alterations of a complementary network property, “fan-out” (the number of unique Purkinje cells innervated by each parallel fiber). To determine if there was an increase in fan-out of parallel fibers to Purkinje cells, at each age we analyzed parallel fiber connectivity to a set of 5-20 consecutively positioned Purkinje cells that were orthogonally oriented to the direction of parallel fibers (Table 1; Fig 2A-E, Fig. S4 A-D and Video 9). We extracted a subset of parallel fibers (144-2155 per age) that were in close proximity (within 5 um) of the contiguous subset of reconstructed Purkinje cells. Among the potential Purkinje cell postsynaptic targets, the percentage of Purkinje cells innervated by a parallel fiber increased steadily during the first two postnatal weeks ranging from a mean of 3.8% at P0 to 25% at P14 (Fig. 2F). This incremental 5.5-fold increase in the synaptogenesis of single axons is relatively minor compared to the ∼500-fold increase we observed in fan-in connectivity of parallel fibers onto Purkinje cells.

**Table 1.**
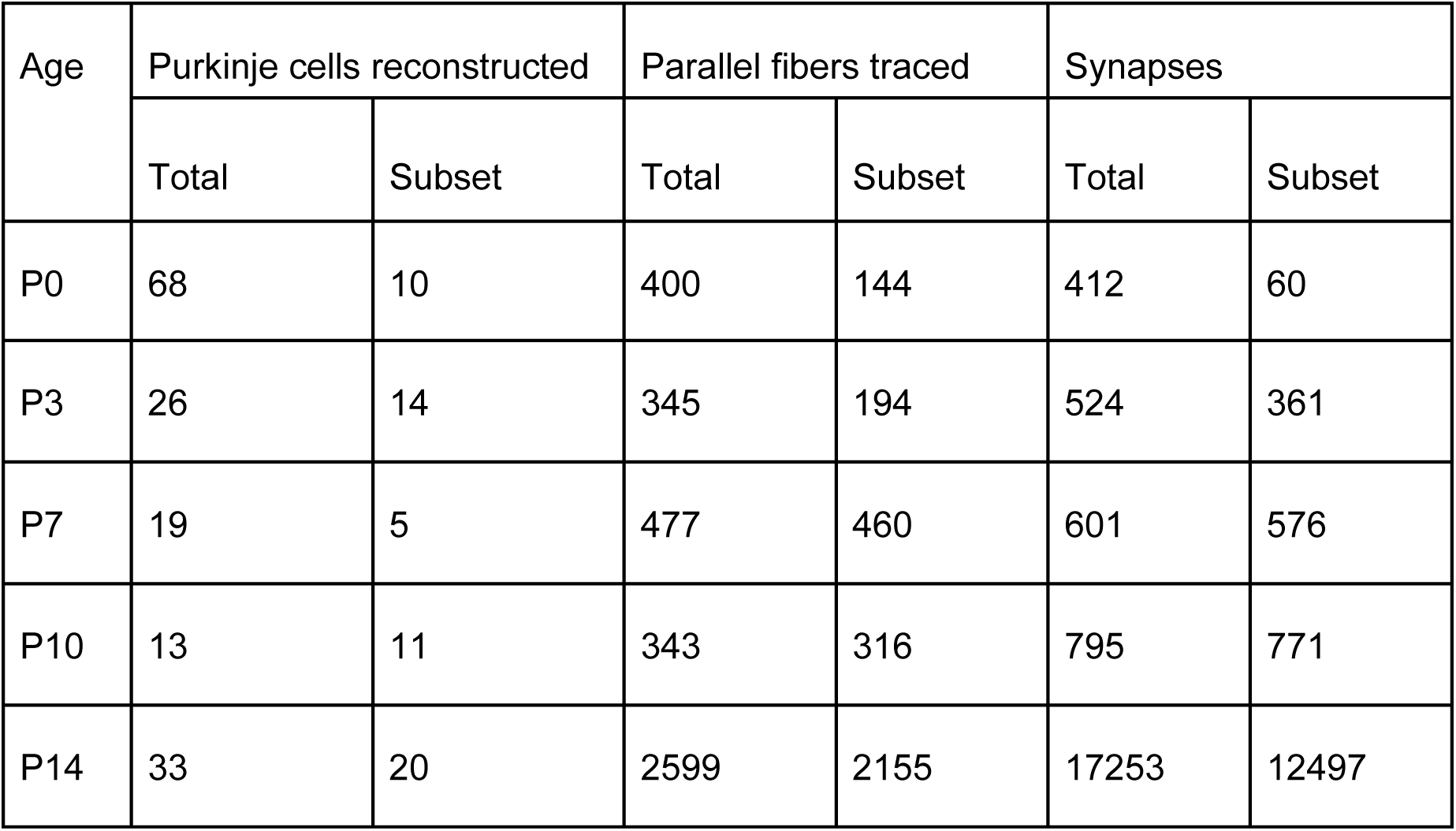
Numbers of parallel fibers and Purkinje cells reconstructed per dataset, and numbers of synapses formed between them. From the total reconstructed dataset, we selected a subset of Purkinje cells and parallel fibers; only those parallel fibers passing within 5 µm of Purkinje cell dendrites were included. All Purkinje cells and parallel fibers within this subset thus had the potential for synaptic connection.

The progressive accretion of connectivity between parallel fibers and postsynaptic Purkinje cells differed considerably from the reorganization of climbing fiber inputs. This means that, over this developmental period, parallel fibers appear to be doing something fundamentally different from climbing fibers, which are concurrently losing connectivity. The climbing fibers appear to eliminate their connections with the majority of Purkinje cells they previously innervated ^54–57^.

### Sequential assembly of mossy fiber-to-granule-cell-to-Purkinje cell connectivity

Interestingly, some parallel fibers that passed through the stack of Purkinje cell dendrites we analyzed (spanning lengths between 50 and 100 µm per age) did not establish synapses with any of them, and we wondered why. One possibility was that granule cells, which extend their parallel fiber axons sagittally for long distances (millimeters or longer), may target Purkinje cells at specific distances, such that parallel fiber connection probability is distance-dependent. If this were the case, then we expected that the connectivity of parallel fibers to their granule cell soma in our datasets might be different from those of parallel fibers that were further away. To test this hypothesis, we selected a cohort of 81 parallel fibers stemming from granule cell bodies in the volume (1 fiber per granule cell) and determined whether they were more or less likely to establish synapses on Purkinje cells than 460 parallel fibers in the same region but with cell bodies outside the volume (and hence further away). Collectively, these parallel fibers overlapped with 5 total Purkinje cells. We observed that 63% (n=51/81) of parallel fibers from in-volume granule cells passed these cells without forming any synapses, whereas significantly fewer (25%, or 116/460) parallel fibers from outside-volume cells passed without forming synapses. Thus, the more proximal portions of parallel fibers had lower connection probabilities than more distal portions.

To explore potential anatomical underpinnings of this reduced connectivity, we examined the granule cell bodies of the 81 parallel fibers above. We found that n=50/81 of granule cells exhibited innervation by mossy fibers, a second type of afferent input to the cerebellum, indicative of advanced developmental maturity. The remaining 31 granule cells had no mossy fiber synapses. Strikingly, the mossy-fiber-innervated group bypassed all Purkinje cells at a frequency of 40% (n=20/50), and the entirety of the mossy-fiber-free group (n=31/31) bypassed all Purkinje cells, suggesting an increased connection probability for more mature granule cells.

We observed evidence of mossy-fiber-related alterations in connection probability at other developmental ages, as well, with the added characteristic that the proportion of mossy-fiber-innervated granule cells increased with developmental age: at P3, 32% (n=19/60) cells were not innervated by mossy fibers, and all of these had parallel fibers that completely bypassed Purkinje cells. Granule cells with mossy fiber innervation at P3, however, bypassed Purkinje cells with 58% frequency (n=24/41; Fig. S4F). At P10, granule cells without mossy fiber input similarly did not form inputs onto Purkinje cells (Fig. S4G). Taking these observations together, we conclude that parallel fiber synapse formation onto Purkinje cells may be predicated upon, or accelerated by, mossy fiber innervation of granule cells.

To explore this idea of a temporal sequence between mossy fiber innervation of granule cells and parallel fiber innervation of Purkinje cells, we took advantage of the orderly topographic layering of granule cells related to their maturational state. As previous studies have shown, the most mature granule cells are located relatively deep in the inner granule cell layer ^47,48,58^. Consistent with this, we found that their dendrites were all innervated by mossy fiber synapses, and all their parallel fibers established synapses with one or typically multiple Purkinje cells that they passed through (red color granule cells in Fig. 2I and the bottom panel in 2H; Video 10). These parallel fiber synapses were on the more proximal dendritic regions of Purkinje cells. The granule cells with somata located slightly more superficially in the inner granule cell layer (i.e. somewhat closer to Purkinje cells) corresponded to parallel fibers that were located more superficially (i.e. somewhat closer to the pia) in the molecular layer. The dendrites of these granule cells were all innervated by mossy fibers. Interestingly however, their parallel fibers did not establish any synapses with Purkinje cell dendrites (green color granule cells in Fig. 2I and middle panel in 2H). Granule cells that were located just below or just above Purkinje cells (i.e. ones that were still migrating radially inward through cerebellar cortex) were not yet innervated by mossy fibers, and their parallel fiber axons did not establish synapses with Purkinje cells (purple color granule cells in Fig. 2I and the top panel in 2H). Their parallel fibers ran through more distal portions of Purkinje cell dendritic arbors. Finally, the least mature granule cells were still located in the outer granule cell layer. Some of these had bipolar morphologies consistent with the migration and/or early differentiation time points in granule cell development ^38,49,50^. Because these cells did not yet have dendrites, they showed no evidence of mossy fiber input (granule cells numbered 1 and 2 in Fig 2G). Their parallel fibers (when identifiable) were located in the most superficial parts of Purkinje cell arbors but did not yet establish any synapses with them. These results argue for a tight association between the maturation of presynaptic input to granule cells and the subsequent postsynaptic output to Purkinje cells. The fact that the least mature granule cells that had mossy fiber input were still unable to generate synapses on Purkinje cells, suggests that mossy fiber input may be required for subsequent establishment of output synapses, and raises the possibility that mossy fiber activation of granule cells may influence the establishment of parallel fiber synapses.

### Purkinje cell filopodia facilitate the assembly of extensive parallel fiber-to-Purkinje cell connectivity

We observed a profound synaptogenesis between many thousands of parallel fibers and each Purkinje cell, beginning at the end of the first postnatal week. A striking phenomenon that coincided with this was the appearance of transient, filopodia-like processes that emanated perpendicularly from Purkinje cell dendrites, which were especially prevalent at the start of this accelerated growth. (Figs. 3A,B). We found that many of these “filopodia” were actually synaptic targets of parallel fiber axon terminals (Fig. 3D). These newly formed synapses were variable in terms of volume and number of synaptic vesicles, both conventional indicators of neurotransmission efficacy (see below). We examined 8,355 filopodia emanating from the 147 dendrites of eight nearby P7 Purkinje cells (Fig S5 A-E), and identified 12,793 synapses on these filopodia in total (Video 12). The greater number of synapses than filopodia was explained by the fact that some filopodia (36%) had more than one synapse (Figure S5K).

**Figure 3.**
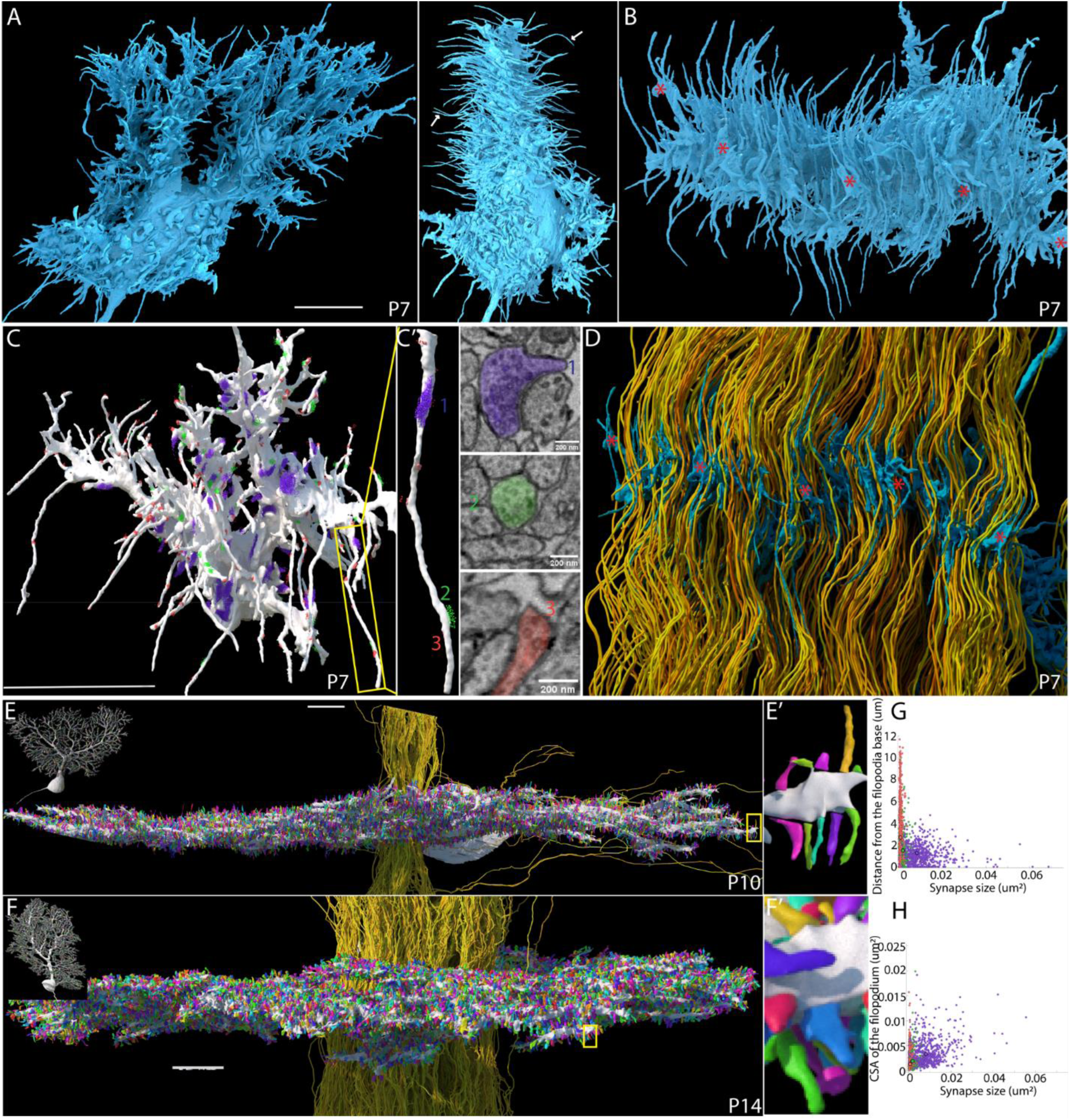
Short-lived developmental stage of profuse Purkinje cell dendritic filopodia oriented in the same direction as parallel fibers. A) A P7 Purkinje cell shown in the sagittal (left) and coronal (right) planes. Purkinje cell filopodia (white arrows) are largely oriented mediolaterally. B) A top-down (radial or dorsoventral) view of the same Purkinje cell showing that its dendrites are becoming oriented in one plane and that the filopodia are oriented orthogonally to that plane. C. These filopodial-like extensions are usually innervated (sites of vesicle clusters in the innervating axons are shown in red, green and purple depending on how many vesicles at the synapse, with purple being the largest vesicle clusters and red the smallest. C’) A single filopodium (the yellow rectangle in panel C) shown at higher power, highlighting three types of synapses. Corresponding electron micrographs illustrating a cross-section of each synapse are shown to its right. D) The same Purkinje cell and view as in B) with the parallel fibers shown in yellow-orange color. These parallel fibers are the source of 85% of the filopodial synaptic sites. E) A P10 Purkinje cell (see inset in top left of panel) viewed from the pial surface. The cell’s dendritic arbor (white) is fully planar. All dendritic spines (18,700) are shown in random colors. Although they have mixed morphologies most appear to be extending in the orientation of the parallel fibers (340 parallel fibers are labeled yellow). E’) A higher magnification of the region in the yellow rectangle of panel E showing that the orientation of the spines is similar to the parallel fibers F) A P14 Purkinje cell rendered the same way as the P10 cell in E. The 18,700 spines have all been labeled with random colors and also show an orientation that is parallel to the Purkinje cells. A P14 Purkinje cell shown from pial view showing spines in the similar orientation to P10 and P14 (color) and in the orientation of the 2599 reconstructed parallel fibers (yellow). F’) A higher magnification portion of the dendrite marked in yellow rectangle showing that the spines remain oriented in the same direction as parallel fibers. G) A swarm plot showing association of synapses along filopodia (distance from base of filopodia), mature synapses (purple circles) are located most proximally to the dendrite (1.25µm, std 0.821). Immature synapses are located closer to the base of the filopodium (1.48, std 1.29 µm) and while developing (nascent) synapses are present all along the length of filopodia distal portions of filopodia contain largely developing synapses (2.68µm, std 2.12). H) A swarm plot showing the association of synapses with the diameter of the filopodia at their respective locations. Mature synapses are present where filopodial diameters are larger (mean 0.0033, std 0.00224 µm^2^), immature synapses are associated with intermediate filopodial cross sectional area (mean 0.0019, std 0.00168 µm^2^), and developing synapses are located at thinner portions of filopodia (0.00132, std 0.00129 µm^2^). Scale bars: 10 µm.

The synapses on the filopodia showed diversity in their structure (Fig. 3C; Fig. S5 H, I; Fig. S5; Video 12) that we were able to separate into three general categories (Fig. S4I): the most mature-appearing synapses (“Type 1”, about 28% of all filopodial synapses) were large (mean volume = 0.1574±0.0051 μm^2^) and had between 50 and >1,000 synaptic vesicles (Fig. S6A). They had prominent postsynaptic densities characteristic of mature, glutamatergic parallel fiber synapses (Fig. S5B; ^43,45^), with wider-caliber postsynaptic filopodial sites (mean ± standard deviation cross-sectional area [CSA] 0.0033 ± 0.00224 µm2). There were also more immature-appearing synapses (“Type 2”, 14%) with 10-50 synaptic vesicles and smaller volumes (0.0434 ±0.0016 µm^2^). These synapses did not show postsynaptic densities and the filopodial caliber at their site of contact tended to be of smaller caliber (0.0019 ± 0.00168 µm^2^ CSA.

The remaining 58% of synapses (“Type 3”) were quite small (0.0184±0.0003 µm^2^) with only 1-10 synaptic vesicles, also lacked postsynaptic densities, and had smaller postsynaptic areas (CSA, 0.00132 ± 0.00129 µm2). We also observed that the location of these three synapse types differed: on filopodia that received more than one parallel fiber synapse, the Type 1 mature synapses were, on average, closest to the dendrite (mean ± standard deviation 1.25 ± 0.821 µm from the parent dendrite), Type 2 synapses were further out on the filopodia (1.48 ± 1.29 µm), and Type 3 synapses were most distal on average (2.68 ± 2.12 µm; Fig. 3G). We identified the same trend on filopodia receiving only one parallel fiber synapse.

The P7 Purkinje cells we examined possessed 900-1900 dendritic filopodia total, of which ∼87% were innervated by parallel fibers (Video 13). All told, ∼74% of the total synapses innervating P7 Purkinje cells were located on filopodia (with 20% formed onto dendritic shafts and the remaining 6% formed onto soma). Notably, although we identified filopodia on Purkinje cells at P3, we observed no filopodial synapses at that age (Fig 1B inset). In combination with data from other ages, this analysis suggests a role of filopodia in generating the rapid rise in parallel fiber to Purkinje cell connectivity: the parallel fiber preference for filopodia represented a switch from strict parallel fiber innervation of Purkinje cell dendritic shafts at P3, and at P10 and P14 all parallel fiber synapses were on dendritic spines. These data suggest that filopodia serve as a transitory component in the process of parallel fiber synapse localization.

### Morphological differences distinguish immature vs. mature parallel-fiber-Purkinje-cell synapses

To better understand how synapses transition to their mature state, we compared immature and mature synapses in two different ways. First, we studied parallel fiber synapses at one part of the dendritic tree at different developmental ages. Second, we studied parallel fiber synapses established in different parts of the Purkinje cell dendritic tree at one age. We reached the same conclusion from both datasets. Our quantification revealed that immature parallel fiber to Purkinje cell synapses have smaller contact area between the pre- and postsynaptic cells when we compared either mature proximal dendritic synapses to immature distal ones at a single age (Fig. S5P and Q) or average contact areas of spine synapses in immature (P7) versus mature (P14) datasets (Fig. S5N). Our spine analysis also revealed that spine metrics became significantly more uniform in older cerebella, indicating a profound reduction in variance. This increased uniformity in filopodial length and volume from P7 to P14 (Fig. S5L and M) is consistent with previous reports that the mouse cerebellar cortex begins to approach full maturity between P14 and P21.

### Parallel fiber input is largely individualized to each Purkinje cell

To determine if there was any pattern in the parallel fiber-to-Purkinje cell connectivity at P14, we examined the connections between a cohort of parallel fibers and Purkinje cells (2155 PFs and 20 PCs) that, as previously explained, were ideally positioned to form synaptic connections (Fig 4A). As shown in Fig S7A, the distribution of parallel fiber fan out (number of Purkinje cells innervated by each) was unimodal and approximately symmetric, with mean 5.02 and range 0 to 12. The distribution of parallel fiber fan-in (number of parallel fibers innervating each Purkinje cell), however, was less regular, appearing to have one mode near 200 parallel fibers/Purkinje cell and a right-tailed distribution centered at a second mode near 500 (total distribution mean 541 and range 200 to 1033; Fig. S7B). To learn if there were some group preferences among parallel fibers for connecting to particular Purkinje cells, we compared the innervation of random PC pairs to assess whether the cohort of parallel fibers that innervated one Purkinje cell was, as a group, more or less likely than chance to innervate another arbitrary Purkinje cell. We found that on average 24.3% of the cohort of parallel fibers innervating each Purkinje cell established synapses on another arbitrary Purkinje cell, which was consistent with this being a chance occurrence given the 25% probability of connectivity between parallel fibers to an individual Purkinje cell mentioned above (Fig. S6C). There was, however, considerable range in this pairwise analysis: at one extreme one Purkinje cell as co-innervated by 45% of the cohort that innervated another Purkinje cell and, at the other extreme, one Purkinje Cell was innervated by only 7.5% of the cohort that innervated another Purkinje cell). To determine the likelihood of this connectivity pattern occurring by chance, we performed a correlation analysis on the shared parallel fibers among the 20 Purkinje cells (see Methods for details). Our analysis demonstrated that the connectivity pattern observed was significantly different (p=0.001) from what would have been expected if connectivity were generated by chance (Fig 4B). The strongest conclusion was that there was less sharing by Purkinje cells than would be expected by random chance. This tendency for non-random connectivity was also supported by other statistical methods that yielded similar findings to the correlation analysis (see Figs. S7E, S8, S9). In summary, our analysis shows that Purkinje cells share fewer parallel fibers than would be expected by chance.

**Figure 4.**
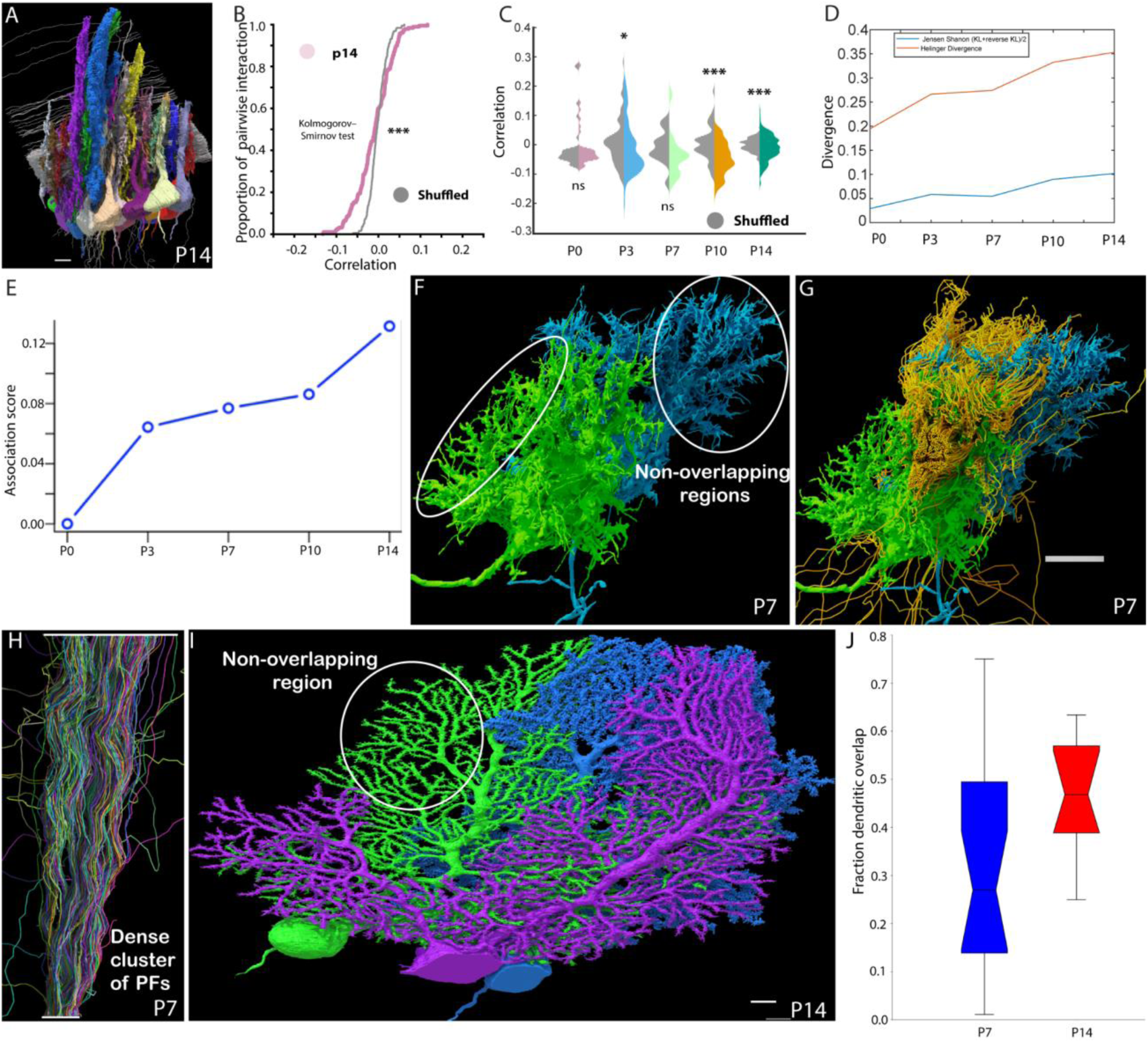
The connectivity of parallel fibers and Purkinje cells is unlikely to have happened by chance. A) Shown is a cohort of 20 adjacent Purkinje cells (in different colors) and 2599 parallel fibers (gray) from a P14 mouse. In the analysis in panels B, C, and D, only parallel fibers that passed within 5 ums of the Purkinje cell dendritic branches of all 20 cells were included. B) A cumulative distribution function plot showing the correlations in the connectivity among pairs (n=190) of these Purkinje cells. A correlation of 0.0 indicates that the number of shared parallel fibers between two Purkinje cells is exactly what would be expected by chance. The actual data is skewed towards anticorrelation, with approximately 70% of pairs showing more differences in connectivity than expected by chance. Conversely, about 30% of pairs exhibit a more positive correlation. The experimental data (pink) is notably more skewed than the distribution derived from shuffling the parallel fiber connectivity (gray curve) C) A similar analysis performed on sets of adjacent Purkinje cells at P0, P3, P7 and P10 and plotted along with P14 (shuffled are shaded gray and colors represent the experimental data) shows that this non-random connectivity is most evident at the later stages of development (P10 and P14). D) The difference between shuffled data and observed data was quantified by the Jensen Shannon divergence (blue) and the Hellinger divergence (orange) from P0-P14. Both analyses show that the connectivity diverges from chance with age. E) Chart of the Purkinje cell-parallel fiber association score across five data matrices, arranged in time order. The increasing association score (see Methods) indicates a growing degree of structured dependence between Purkinje cells and parallel fibers over time. F) Two adjacent P7 Purkinje cells with partially non-overlapping dendritic trees. G) The same two Purkinje cells shown with a set of ∼500 parallel fibers passing through the region where the arbors of the dendrites of two Purkinje cells overlap. Given the density of these axons, there are likely even more parallel fibers that are not shared. This also implies that the parallel fiber connectivity to even adjacent Purkinje cells will be significantly different. H) A cluster of adjacent parallel fibers were traced for 75 µms in a P7 mouse. The results show that these parallel fibers are more divergent than parallel with a spread of ∼2.5-fold over 75 ums. I) Non-overlapping regions of adjacent Purkinje cells were also evident at P14. Shown are 3 Purkinje cells labeled blue, green, and purple. One of the non-overlapping regions is shown within the yellow circle. J) The fraction of dendritic overlap between any two Purkinje cells is on average less than 50%, meaning that the parallel fiber connectivity of each Purkinje cell is individualized. Scale bars: 10 µm.

The less-than-expected amount of sharing of parallel fibers between Purkinje cells at P14 raised the question of how this pattern might have emerged. One possibility was that a greater amount of sharing was present at earlier times but after a stage of parallel fiber pruning, their connectivity became less paired. Alternatively, Purkinje cells might have shared fewer parallel fibers than expected by chance throughout development and the large degree of synapse addition was selective to maintain this non-random distribution. To investigate the likelihood of these options, we performed similar analyses on the four earlier developmental stages: P0, 3, 7, and 10. We found that from P10 onwards, Purkinje cells shared fewer parallel fibers than expected by chance (based on an entropy test; Fig 4C, Fig. S7F-I and Fig. S8, p<0.001; Kalmogorov-Smirnov’s test, Methods). Interestingly, as the overall connectivity between parallel fibers and Purkinje cells increased, the deviation between the observed connectivity patterns and the random simulations became more pronounced, suggesting that newly added synapses were not formed purely by chance (Figs. 4D, E) giving each Purkinje cell somewhat unique cohorts of parallel fiber input.

## Discussion

We have analyzed the dendritic geometry of Purkinje cells and their parallel fiber inputs in developing mice at five time points between birth and 2 weeks of age, and our findings have revealed various aspects of a complex, dynamic maturation process in this system. Notably, Purkinje cells exhibit dramatic changes in shape and size over the first two postnatal weeks while the parallel fiber to Purkinje cell connectivity increases 500-fold, beginning with a slow phase during the first week following birth and accelerating substantially between P7 and P10. This timing overlaps with the critical window for climbing fiber input selection onto Purkinje cells: light-microscopy-based studies of climbing-fiber-to-Purkinje-cell synaptic development indicate P7-P10 as a key timeframe for climbing fiber competition and pruning, with a single climbing fiber maturing onto each Purkinje cell from P10 onward ^59^. Our observation of this overlap is interesting in light of more recent empirical models of cerebellar function, which indicate that at mature Purkinje cells, parallel fibers input modules encode input “features” and the singular climbing fiber input is a corresponding error signal. Given that parallel fiber and climbing fiber selection processes are known to be activity-dependent ^60–64^, and that the resulting inputs are tuned to the same input signals (e.g. sensory or motor), our finding that both processes resolve at nearly the same time suggests that they are closely interdependent on each other. Throughout early postnatal development, we observed that granule cells do not form output synapses (via parallel fibers) onto Purkinje cells until they first receive input from mossy fibers (which carry the aforementioned input signals). This supports a model in which mossy fiber input serves as a critical gatekeeper for granule cell integration into the cerebellar network, representing an example of sequential circuit assembly: afferent reception precedes and possibly instructs efferent output formation. Such a sequence provides a mechanism for ensuring that only granule cells already incorporated into information-processing streams become functionally engaged with Purkinje cells.

Corresponding with the period of rapid circuit expansion and reorganization, we observed that parallel fiber synaptogenesis during the second week seems to be mediated by filopodia. One possible utility of these filopodia might be to enable flexibility during parallel fiber synapse formation, since many parallel fibers are already in place (having grown mediolaterally) by the time postnatal synaptic remodeling begins, and since functionally preferable parallel fibers may be further away from the Purkinje cell than its young dendritic processes. Interestingly, these filopodia, despite forming synapses with parallel fibers, also overwhelmingly ran in parallel with the parallel fibers, a characteristic not typical of axon-to-dendrite connections in other mammalian brain regions ^65,66^. Another functionality of these filopodia may thus be to facilitate proper Purkinje cell orientation during the first week. Additional studies of these compartments should be undertaken to clarify their role in cerebellar development.

The comprehensive connectivity analysis showed less sharing of the same parallel fibers on Purkinje cells in our datasets than expected by random chance. This was the case despite our correcting for anatomical characteristics that might reduce sharing between Purkinje cells, including oblique tiling of Purkinje cell dendritic arbors (such that only fractions of dendritic arbors between consecutive Purkinje cells overlapped; at P14 we measured 50% overlap; see Figs. 4F, G, I, and J) and large-spatial-scale shifts in parallel fibers away from truly mediolateral trajectories. Our results thus indicate the presence of a factor aside from these cytoarchitectural elements (which may arise due to genetic signaling or activity-based signaling) that results in individualized the input to each PC. This establishment of unique parallel fiber cohorts onto Purkinje cells in a localized cortical region may further suggest a dependence on climbing fiber signaling, since in a previous study we also found that a unique climbing fiber had established prominent synaptic connectivity with each Purkinje cell in a localized region by P7 ^67^.

### Principles of granule cell wiring

Our analysis revealed several principles related to how parallel fibers become connected to Purkinje cells. First, we found that parallel fibers predominantly innervated individual Purkinje cells with one synapse at all the five developmental stages analyzed. At each age, between 80% and 90% of the parallel fibers established one synapse on the Purkinje cells they innervated (Fig. S2V). This is also the case in most other parts of the mammalian cerebrum where axons incidentally innervate dendrites along their relatively straight paths ^68–71^. Second, we find evidence that at P14, a minority of parallel fiber axons (30%, or roughly 36,000 of 120,000) pass through a PC (within 5 µm of the dendrites) actually form synapses. This is interesting in the context of another recent connectomic study in cerebellum performed at an older developmental age, which reported that 50% of parallel fibers passed Purkinje cells without forming synapses ^41^: both these studies contrast with previous estimates of nearly 100% parallel fiber to Purkinje cell innervation frequency (in adult cat cerebellar cortex; ^72^). This discrepancy may reflect either the better resolution of EM-based connectomic studies, a difference in parallel fiber remodeling dynamics across mammalian species, or possibly an additional, later phase of parallel fiber remodeling. Further studies will be useful for examining these possibilities. Third, we found that during the first two postnatal weeks, each parallel fiber increases the number of Purkinje cells it innervates. This steady and progressive additional of connectivity (2% at P0; 12% at P3; 17% at P7; 22% at P10; and 30% at p14) is unlike the eliminative process going on for certain other developing axons (e.g. climbing fibers in the cerebellum ^54–57,67,73^, sympathetic and parasympathetic axons in the autonomic nervous system ^74,75^ or motor axons in skeletal muscle ^22,76,77^.

In summary, we have used comparative connectomics to assay connectivity in the developing mouse cerebellum, analyzing synapse-level data from cerebella between birth and postnatal day 14. Our findings indicate a highly dynamic process that entails multiple phases of parallel fiber synapse addition, resulting in hundreds-fold additional synapses by the end of the first 2 weeks of postnatal life. We further explore characteristics of filopodia that may appear to facilitate this addition. Finally, we report observations that indicate a close relationship between parallel fiber and climbing fiber input selection (near-coincident timing and evidence of parallel fiber/climbing fiber cohort uniqueness among nearby Purkinje cells) and a potential role for mossy fibers in mediating this process. These insights may serve as helpful points of focus for future studies of both normal and disordered development in the cerebellum ^78–83^.

## Materials and methods

### Animals

Mice were maintained on a 12 h:12 h light–dark cycle, with ad libitum access to food and water, in a temperature - (22 °C) and humidity - (30–70%) controlled environment. All experiments were performed in accordance with NIH guidelines and were approved by the Harvard University Institutional Animal Care and Use Committee (IACUC).

### Electron microscopy

Mice were anesthetized with isoflurane and positioned ventral side up. The chest was opened, and a needle was inserted through the left ventricle into the ascending aorta. Using a peristaltic pump, transcardial perfusion was performed at a flow rate of 10-15 mL/min, beginning with fresh aCSF for 2-3 minutes to clear the blood. Following this, an ice-cold mixture of fixatives (2% glutaraldehyde + 2% PFA in 0.15 M sodium cacodylate) was perfused for at least 3 minutes to fix the brain.

Once the brain was removed after opening the skull, it was placed in the same fixative and kept on a slow shaker overnight in a cold room. The fixed brain was then sectioned into slices of the desired thickness (200-750) using a vibratome, and these slices were fixed overnight in the same fixative. After fixation, the slices were washed three times for 10 minutes each with sodium cacodylate buffer.

Next, 2% OsO4 was added to the slices, which were positioned in the center of a thick spacer (200 um, Invitrogen, Thermo Scientific, S24737), on a glass slide and covered with another coverslip to prevent deformation. After 30 minutes, the slices were transferred to mini vials containing 2% OsO4 solution for 4 hours on a shaker, followed by three washes for 10 minutes each with sodium cacodylate buffer to ensure the removal of any OsO4 traces.

Freshly prepared 1% K4Fe(CN)6 was added, and the vial was placed on a shaker for 2 hours at room temperature, allowing the solution to turn bluish green. After this, filtered 1% thiocarbohydrazide (TCH) solution, prepared in deionized water (TCH was dissolved by heating it at 45°C), was added, and the slices were incubated for 1 hour, with heat or sonication used to dissolve TCH. The slices were washed again three times for 10 minutes each with deionized water.

After the TCH treatment, an additional osmium tetroxide treatment, 2% OsO4, was added, and samples were kept on a rotator or shaker for 4 hours. Following this, the slices underwent three more washes for 10 minutes each with distilled water. They were then treated with a 2% aqueous uranyl acetate solution, stored at 4°C overnight, and later moved to room temperature for 15 minutes before being placed in an oven at 60°C for 1 hour. After cooling to room temperature, the slices were washed three times for 10 minutes each with distilled water.

To dehydrate the slices, a graduated ethanol: water series was employed: 30% alcohol for 10 minutes, followed by 50%, 75%, and 95% alcohol for 15 minutes each, with a final three changes of 100% alcohol. After rinsing the slices twice for 5 minutes each with 100% propylene oxide or acetone, they were infiltrated with a thoroughly mixed combination of propylene oxide or acetone with resin (Epon 812 or LX112). 25% resin in propylene oxide mix was added for 30 minutes, then 50% and 75% resin in propylene oxide for 6-12 hours each, and finally 100% resin for 12 hours each. Finally, the slices were placed in the desired orientation within molds and cured at 60°C for 48-72 hours.

### Cutting serial sections

30 nm serial sections were cut and collected using ATUM (Automatic Tape-collecting Ultra-Microtome), as described in Kasthuri, et al., 2015 ^84,85^. Briefly, the resin block face was trimmed with a Diatome trimming knife. Then serial sections (1800 for P10 and ∼8900 for P14) of 30 nm were cut using Diatome diamond knives (35 degree or 45 degree) on Leica ultramicrotome with ATUM set up attached for automated collection of the sections on Kapton tape.

#### Wafer making

Circular silicon wafers (<100>) were cut into rectangles to be compatible with multiSEM imaging. Black double-sided carbon tape (Ted Pella, 16084-8) was then laid onto the rectangular silicon wafer. The Kapton tape with sections was then cut into pieces of length similar to a silicon wafer and laid on top of double-sided carbon tape.

#### Post staining

2% aqueous Uranyl acetate solution was filtered and poured onto sections on silicon wafers. The solution was washed off after 5 minutes using distilled water 3 times, and the wafer was dried using forced air. A similar protocol was performed for 1% lead citrate (Leica Ultrostain II, 16705530). The wafers were stored in vacuum until imaging.

#### Wafer mapping

Wafer images were captured using the Zeiss Axio imager microscope. The boundaries of serial sections were defined in the wafer images using the Serial Array Tomography (SAT) program in the mSEM Zeiss Zen imaging software. The region of interest (ROI) to be imaged in the SEM was mapped using the same program. The mapping was then transferred to the multi-SEM computer for imaging. Using the SAT program the position of the sections was then aligned for the automatic stage movements and imaging of all the sections in the wafer.

#### multiSEM imaging

Each multiSEM (Zeiss) field of view (mFoV) covers an area of ∼100 um in width. Each mFoV consists of 61 tiles imaged by the 61 beams. The sections were imaged at 4×4 nm resolution using a dwell time of 200-800 nanoseconds per pixel. The microscope operated at a voltage of 1.5 kV with an individual beam current of 575 pA and a tile overlap of 1%. Image quality (signal to noise, focus, and other artifacts) was checked in real time, and problematic sections were reimaged at the end of each image cycle. The images were then stitched and aligned.

### Stitching and Alignment

Rigid stitching was performed on most of the raw image tiles coming from the microscope. For some more distorted images we applied affine stitching. Firstly, scale-invariant feature transform (SIFT) features were extracted from the boundary area of each raw tile. Then all the neighborhood tiles in each section were matched together based on SIFT feature matching. Finally, a global optimization step made the matching results smooth. The optimization step differed for rigid vs. affine stitching.

After each section was stitched, an elastic alignment of all the sections was performed. A rough affine transformation between each mFoV in section i to section i+1 and i+2 was estimated based on the SIFT features of blob-like objects detected in the sections. Then template matching which performed as a fine-grained matching for section i to section i+1 and i+2 was done on grid distributed image blocks in section i. Finally, a global optimization on all the sections in the whole stack made the elastic transformation of mesh grid points on all the sections smooth.

The aligned stack was rendered at full resolution, and each section was cut into 4096*4096 *.png tiles for further analysis.

### Neuronal reconstruction in P0, P10 and P14

Aligned EM stacks (voxel size 4×4×30 nm³) were reconstructed with a semiautomated workflow that first generates machine-learning 2D membrane segmentations and then uses those predictions to guide 3D proofreading and reconstruction. Membrane-probability maps were produced on selected 3D portions of the volumes with a pretrained 2D U-Net ^86^ run through mEMbrain ^87^, an interactive MATLAB front-end for connectomic segmentation that interfaces with VAST ^88^. In VAST, the selected regions perform model corrections. mEMbrain then sampled 256 × 256-pixel patches at random positions and arbitrary orientations, but only from voxels that already carried a ground-truth label; this random sampling itself provided the flips and rotations that serve as data augmentation. After each training round, the annotator opened newly chosen regions anywhere in the aligned volume within VAST, viewed them in full ultrastructural context and directly corrected the previous-round network predictions. The updated labels were used to fine-tune the network for 1-4 epochs per round, each epoch presenting 20,000-100,000 augmented patches, until validation accuracy plateaued. Three such cycles, representing approximately 70 h of annotation, produced the final model.

For section-wise instance segmentation, the probability maps were down-sampled to 8 nm in-plane resolution with bicubic interpolation. Local minima of the h-minima transform of the down-sampled map ^89^ served as seeds, and mEMbrain’s watershed-based region growing yielded non-overlapping 2D object masks. During proofreading and reconstruction in VAST the probability and segmentation layers constrained brush strokes: ML scores limited edits to high-confidence voxels, and the current object ID restricted edits to a single planar object. These constraints accelerated semi-automatic 3D agglomeration and produced high-quality segmentations with less manual effort ^87,90^. All trained networks generated in this study are provided to the community.

### 3D neurite segmentation of P3 and P7 cerebella

We utilized the PyTC software package ^91^ for both neuron segmentation and synapse detection.

For **neuron segmentation**, we began by densely annotating neural structures in a subvolume of the P7 dataset, measuring 15μm×15μm×1.8μm. This subvolume included representative structures such as parallel fibers, granule cells, and Purkinje cells. Using these annotations, we trained a 3D UNet model ^86^ to predict the 3D affinity map for each voxel. During inference, the trained model was applied to the 3D image volume, and the predicted affinities were converted into instance segmentations using the Water package ^92^. However, due to imaging noise and volume misalignment, many parallel fibers were erroneously split into multiple instances. To address this, we employed an Intersection-over-Union (IoU)-based tracking method to detect and correct false split errors.

For **synapse detection**, we initially used a pretrained synapse detection model ^93^ via the PyTC software to identify the pre- and postsynaptic components of each synapse. Subsequently, we manually proofread a subvolume of the P7 dataset measuring 13μm×13μm×6μm. The proofread data was then used to fine-tune the synapse detection model. The fine-tuned model was applied to the entire dataset to improve the accuracy of synapse prediction.

### Identification of Purkinje cells during the first postnatal week in mice

Purkinje cells in P0 and P3 cerebella were recognized based on their size compared to other cell types in the cerebellar cortex, location ^49,94^ and based on their high mitochondrial content compared to other cell types. The molecular layer interneurons (MLIs) being inhibitory in nature could potentially complicate the unambiguous identification of Purkinje cells. However, it has been shown that the molecular layer interneurons arrive in the cerebellar cortex later than P3 ^95^. In addition, at P3 and P7, the climbing and mossy fiber innervation of putative Purkinje cells also helps identifying Purkinje cells ^67^, as these afferent fibers are not known to innervate the inhibitory interneurons. In addition, their high mitochondrial content (Fig. S2S), extensive parallel fiber innervation, the beginning of MLIs innervation and the almost monolayer organization of Purkinje cell somata (Fig. 2C) are consistent with the characteristics of Purkinje cells.

### Proofreading and manual segmentation

Neuronal segments generated through automated segmentation methods were imported into the VAST (volume annotation and segmentation tool) ^88^ program for proofreading and for additional manual volumetric segmentation of Purkinje cells, granule cells and their synapses. Dense clusters of axons at all ages were traced in VAST using “skeletonization”, manual definition of a center line using nodes and edges. This method was used for parallel fibers, climbing fibers, and mossy fibers, except for axons connecting with a Purkinje cell at P0 and P7 which were instead volumetrically segmented based on automatic segmentation.

### Tracing dense clusters of parallel fibers and their synapses

Dense clusters of 500 - 4000 axons, at all the five developmental stages, were manually skeletonized in VAST by placing nodes every 10-20 sections at the center of the axon. These nodes are connected by edges forming a continuous skeleton line throughout the length of the axon. The synapses on the axon were annotated by labeling synapse nodes with the name of the postsynaptic neuron. This kind of tracing allows for the generation of a connectivity matrix between axons and their postsynaptic Purkinje cells.

### Quantification of parallel fiber to Purkinje cell connectivity

Skeleton data generated as described above was either saved to .CSV files and imported into MATLAB (The Mathworks, Inc.), or Matlab accessed skeleton data directly from VAST via the VAST API. In MATLAB, connectivity statistics were extracted based on the annotations, and coplot bar graphs and histogram plots were generated.

### Analysis of filopodia and spines

Filopodia and spines (from here on called ‘spines’) were individually segmented (painted volumetrically by hand) in selected Purkinje cells using VAST. In P7, 8 Purkinje cells were completely segmented with 8,355 spines/filopodia total. For P10 and P14, a fraction of proximal and distal spines on the dendritic tree of one cell each were selected and analyzed separately (1000 proximal and 1020 distal spines in P10, and 860 proximal and 730 distal spines in P14). MATLAB was used to extract the physical features of length, volume, and synapse area. The MATLAB script used the VAST API to load and process voxel data for each spine. Spine volume was computed by counting voxels. Lengths were extracted by skeletonization of the volumetric segmentation. The skeletonization algorithm assumes that the spine is a longitudinal branch-free object. It uses filling (seeded 3D flood filling) from a seed point which is located towards the base of the spine and produces a distance field from the seed point, counting steps of filling neighbor voxels. The furthest filled voxel then marks the distal end of the spine. Filling back from the distal-most voxel using the same algorithm identifies the other end. Averaging coordinates of voxels with the same fill distance, at regular intervals, produces locations for nodes of a center-line skeleton which is then copied back into VAST for visual inspection. The length of the skeleton line provides an estimate of the spine length.

Synapses were individually and manually segmented as well, in a separate data layer from spines. Individual synapses were assigned to individual spines by evaluating volumetric overlap of voxels. Since all synapses were painted with the same pen width, synapse area could be estimated by dividing synapse segment volume (based on voxel count) by pen width. In P7, three different maturity levels of synapses were distinguished based on EM appearance (number of vesicles in the axonal bouton, contact area of synapse, and presence or absence of postsynaptic density) and analyzed separately. For mature synapses the electron dense area under the postsynaptic membrane is labeled and measured and this electron dense feature is usually absent in the immature and developing synapses. For all the three classes of synapses vesicle number is counted and the contact area of pre- and postsynaptic neurites is used as additional measures to classify synapses.

Results were plotted using MATLAB. Code is provided in the resources (analyzespines_p14_p10_p7.m).

### Extraction of vesicle numbers

For dendrite 1 of Purkinje cell PC2 in the P7 dataset, vesicles were individually painted in VAST, using a different segment ID per vesicle. All vesicles of a synapse were grouped in a folder labeled by the synapse name. The vesicle count for a synapse is then just the number of vesicle IDs in its vesicle folder, excluding unused IDs. Additionally, vesicle folders were sorted into three different superfolders depending on synapse type 1, 2 or 3. Vesicle folders for each synapse were associated with the corresponding pre/postsynaptic synapse labels by using matching segment names. This analysis is implemented in the Matlab script ‘nag_syn_vesicle_analysis.m’.

### Skeletonization of P7 parallel fibers from volumetric segmentation

In the P7 data, parallel fibers were originally reconstructed volumetrically in VAST using voxel painting. To make downstream analysis consistent with the other datasets, we automatically skeletonized them using a MATLAB script. First, the volumetric segmentation of parallel fibers was loaded into MATLAB. We then computed bounding boxes for all parallel fibers and checked for gaps in the segmentation by using connected component analysis. We attempted to close gaps automatically by using imdilate/imerode. Connected regions were then skeletonized using a seeded filling algorithm as described above (Analysis of filopodia and spines’) and skeletons were uploaded to VAST. Remaining gaps were then corrected by manually connecting any disconnected skeleton parts of a parallel fiber in VAST. Code is provided in the resources (skeletonize_p7_pfs.m).

### Image rendering

All 3D images of cells and other objects in the main figures were rendered using 3ds Max (Autodesk Inc.). Surface meshes of volumetrically segmented objects were extracted from VAST using the MATLAB script vasttools.m. Raw surface meshes were smoothed using ZBrush (Pixologic Inc. / Maxon Inc.) and imported into 3ds Max. Vesicles were modeled by instancing sphere models at vesicle locations using vasttools.m. Skeletons were also exported from VAST using vasttools.m, loaded into 3ds Max and fleshed out to cylinders with a radius of 0.1 micrometers using the ‘Sweep’ modifier. Scenes were illuminated with a textured skydome and rendered using the Arnold renderer.

Some panels in the supplementary figures were exported directly from the VAST 3D viewer.

### Analysis of connectivity patterns

A subset of parallel fibers that were in synaptic reach with a cohort of Purkinje cells were isolated using a program called “convex hull”. To find a subset of axons and Purkinje cells that are fully connected (meaning each axon in the subset has a chance to make synapses on each of the Purkinje cell in the subset, and each Purkinje in the subset has a chance to have synapses made from each of the axon in the subset), we generated a convex hull based on the 3d voxels of each Purkinje cell. Any axons which fall in the convex hull will be considered as ‘has a chance to make synapses on this Purkinje cell’. We exported the tracing/segmentation result of each Purkinje cell from VAST; each section is a single .png file, with the pixels belonging to each Purkinje cell coded with different colors. After extracting pixels of each Purkinje cell in each section, we have the 3d voxels for each Purkinje cell. Then, we dilated the 3D voxels representing each Purkinje cell by 3 μm to make the outer surface smooth. And when generating convex hulls for each Purkinje cell using alphaShape function in MATLAB, we set the smallest alpha radius that produces an alpha shape that encloses all points to 5 μm, because we believe that if an axon is within 5 μm from a Purkinje cell, it is possible to make a synapse on the Purkinje cell. Then for each axon we could calculate whether it’s within the convex hull of each Purkinje cell and we saved it as a m*n 0-1 mask matrix **M** (m axons and n Purkinje cells, **M**ij=1 means axon i is within the coverage of Purkinje cell j, 0 vice versa).

Then finding the largest fully connected subset in the matrix **M** could be considered as a Maximum Clique Problem, which is a Non-deterministic Polynomial Complete (NP-C) problem. We used a greedy strategy to find such a subset.

To assess how similarly different Purkinje cells (PCs) received input from a common set of parallel fibers, we represented each neuron’s input pattern as a binary vector, where each element indicated the presence or absence of input from a specific parallel fiber (PF). We computed pairwise similarities between these input vectors using three complementary metrics: Hamming similarity, Jaccard similarity, and Pearson correlation.

Hamming similarity measures the proportion of input sources for which two neurons exhibit identical input states, including both shared presences and shared absences. This provides a global measure of agreement across all possible inputs, assuming binary encoding. In contrast, Jaccard similarity considers only the subset of sources from which at least one neuron receives input. It quantifies the fraction of these active inputs that are shared between the two neurons, emphasizing functional overlap in received inputs rather than total agreement. Pearson correlation extends the analysis to capture the degree to which two neurons co-vary in the pattern of their inputs across PFs, reflecting the strength and direction of a linear relationship between input vectors.

Together, these metrics provide a multifaceted view of input similarity: Hamming reveals strict pattern matching, Jaccard highlights overlap in active inputs, and correlation assesses coordinated variation in input structure.

To generate the shuffled control (referred to as the configuration model in Fig. 4), we randomly permuted the “connected” status among all PFs connected to a given PC, while preserving the number of inputs per PC. We then recomputed the same three pairwise similarity metrics across the population.

To quantify differences between real and shuffled similarity distributions, we used the Jensen-Shannon divergence (JSD), a symmetric, bounded measure of distributional dissimilarity derived from the Kullback-Leibler divergence. A higher JSD indicates that the real data exhibits structured input similarity not captured by the shuffled model. For statistical comparisons between distributions, we used the Kolmogorov–Smirnov (K–S) test.

### Monte Carlo analysis for assessing shared parallel-fiber connectivity

We tested whether Purkinje cells (PCs) share fewer or more parallel-fiber (PF) inputs than expected by chance with a two-step procedure. Step 1. Pairwise test: for every PC pair we computed, under independent random innervation, the exact probability of observing the recorded number of shared PF axons, given each cell’s total PF-input count and the size of the PF pool. Summing up, the upper and lower tails of this distribution yielded the probabilities of excessive sharing (Pshare) and sharing avoidance (Pavoid), forming complete Pshare and Pavoid matrices for the empirical connectome. Step 2. Network-level benchmark: we generated 1,000 random connectomes using two degree-preserving configuration-model variants; (i) a hard-constraint variant that exactly matches the observed in- and out-degree of every neuron, and (ii) an asymptotic variant in which each potential edge is present with probability p_i = k_i/(N-1), so each node’s degree is freshly drawn from a binomial distribution with mean equal to its observed degree. For every surrogate we recomputed Pshare and Pavoid and extracted two summary statistics: 1) the number of PC pairs with Pshare<0.05 or Pavoid<0.05, and 2) the sum of log-probabilities across all pairs. The Monte Carlo p-value, defined as the fraction of surrogates whose statistic equals or exceeds the experimental value, indicates whether shared PF connectivity can be explained solely by degree structure or instead reflects additional organisational principles (more detailed derivation in ^96^ configuration-model concept ^97^.

### Latent factor analysis using singular value decomposition

For each data matrix X of dimension n by p, where n is the number of parallel fibers and p is the number of Purkinje cells, we apply singular value decomposition to obtain the leading right singular vectors *{u_i_}* and left singular vectors *{v_i_}* of X. Intuitively, for each *^i^*, the right singular vector *u_i_* ɛ *R^n^* characterizes a latent factor among the n fibers, representing a specific Purkinje cell-parallel fiber connection pattern that highlights groups of closely related (co-occurring) parallel fibers; the left singular vector *v_i_* ɛ *R^n^* are the cell loadings indicating the contribution from each Purkinje cell (positive, negative, or zero) to such a latent factor. For each data matrix, the number of informative latent factors is determined using the “eigen gap” criteria suggested in Ding and Ma ^98^. As a result, at most 4 latent factors were chosen across all 5 data matrices, and on average, the leading 4 latent factors can explain at least 65% of the total variations in the data.

### An entropy-based fiber-cell association score

For each data matrix X and its latent factors *{u_i_}*, we propose an entropy score that quantifies the total amount of informative, deterministic patterns that are not likely caused by random chance. To this end, we develop an entropy-based test statistic for the null hypothesis that, for each latent factor, the distribution of its components is of no difference from what would be expected when the fiber-cell connection is completely at random. In particular, we consider the null model where, for given marginal counts of total parallel fiber connections per Purkinje cell (column sums of X), the assignment of the connection counts across n parallel fibers per Purkinje cell follows a multinomial distribution with equal probability 1/n. Specifically, for each latent factor *^i^*, we compute its empirical Shannon entropy using a KNN estimator ^99^ (k=100), denoted as *T_i_(X)*. We simulate the null distribution of *T_i_* by generating N=5000 null matrices of the same dimension as X, whose column sums are identical to X, but the entries are generated based on the null model, a multinomial distribution with equal probability 1/n. Then we compute *T_i_* for each null matrix, generating an empirical null distribution of it. We evaluate to which percentile of the null distribution the based on the actual data X equals. The percentile is denoted as *p_i_*. By definition, a smaller value of *p_i_* (or smaller entropy) indicates a stronger localization pattern in the latent factor, suggesting the existence of groups of co-occurring parallel fibers that are due to intrinsic structural dependence between parallel fiber and Purkinje cells. For each data matrix, we define the parallel fiber-Purkinje cell association score as *E = ∑_1≤i≤4_ log_10_p_i_*, with a higher value indicating a greater amount of structured dependence between the Purkinje cells and parallel fibers. Applying this method to all data matrices, we observe an increasing association score over time, indicating a growing degree of structured dependence between cells and fibers (Fig. 4E).

### Factor analysis of P14

Applying the above methods to P14, we find that the first latent factor indicates the total amount of connected parallel fibers for each Purkinje cell (Fig. S8E), whereas the second latent factor highlights the diversity or distinct groups of fibers connected to different Purkinje cells (Fig. S8F).

### Repressed fiber-sharing among the 18 cells in P14

After removing the two cells in P14 with the highest number of parallel fiber connections, we focus on the remaining 18 cells to examine the extent of co-occurring fiber associations. We analyze the second latent factor, which, as shown in Figure S8G and H, captures parallel fiber groups shared across Purkinje cells. Notably, we observe a higher-than-expected entropy for this factor among the 18 cells (Fig. S8I), suggesting a structured and even distribution of parallel fibers. To quantify this observation, we invert the entropy-based score *p_2_* to construct a one-sided test, using *1-p_2_* as the p-value to assess whether the observed fiber assignment is more uniform than expected under random connections. This yields a p-value of 0 (<2×10^-4^), indicating a significant deviation from the null distribution to the left.

### Quantification of Purkinje cell dendritic overlap

Reconstructed Purkinje cells were exported at 64nm isotropic resolution. Similarly, skeletons of parallel fibers were also exported from VAST. The orientation of parallel fiber trajectories was estimated by averaging the directions of these parallel fibers using X, Y, and Z coordinates in the volumes. The Purkinje cell reconstructions were then projected into 2D images along the average orientation of parallel fibers in the volumes. Overlap between the Purkinje cell 2D projections was quantified using the Intersection over Union (IoU) metric, following dilation of the projected masks by 5 micrometers.

## Acknowledgements

We are grateful to the undergraduate students for their effort in reconstruction of neurons: Daniel Klionky, Emily Yin, Sherryl Deakin, Diqi Zeng, Krishna Swaroop K, Ciara Burke, Kanglin, Zhiwei He, Sneha Aggarwal, Filipa Rodrigues Torrão, Paolo Cucurachi, Wenbo Zhao, Zikun Zhu, Lingdang Zhang. Nagaraju Dhanyasi thanks Juan Carlos Tapia for his valuable discussions on cerebellar development.

---

**Figure S1.**
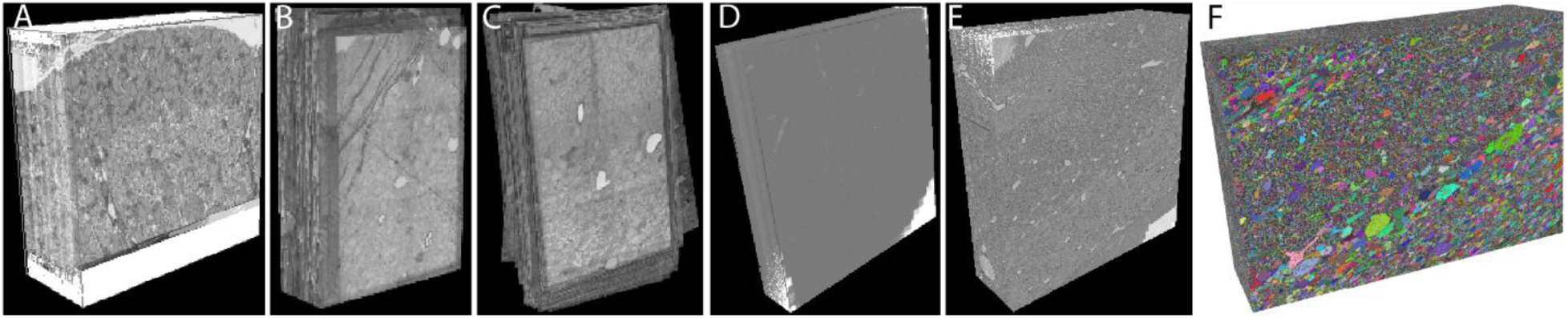
Acquisition and segmentation of electron microscopy images of the developing cerebellar cortex during the first two weeks of postnatal life. A-E) Volumes of cerebellar cortex from P0, 3, 7, 10 and 14, with sizes of the volumes. F) Segmentation of neurons in the P14 volume using machine-learning algorithms.

**Figure S2.**
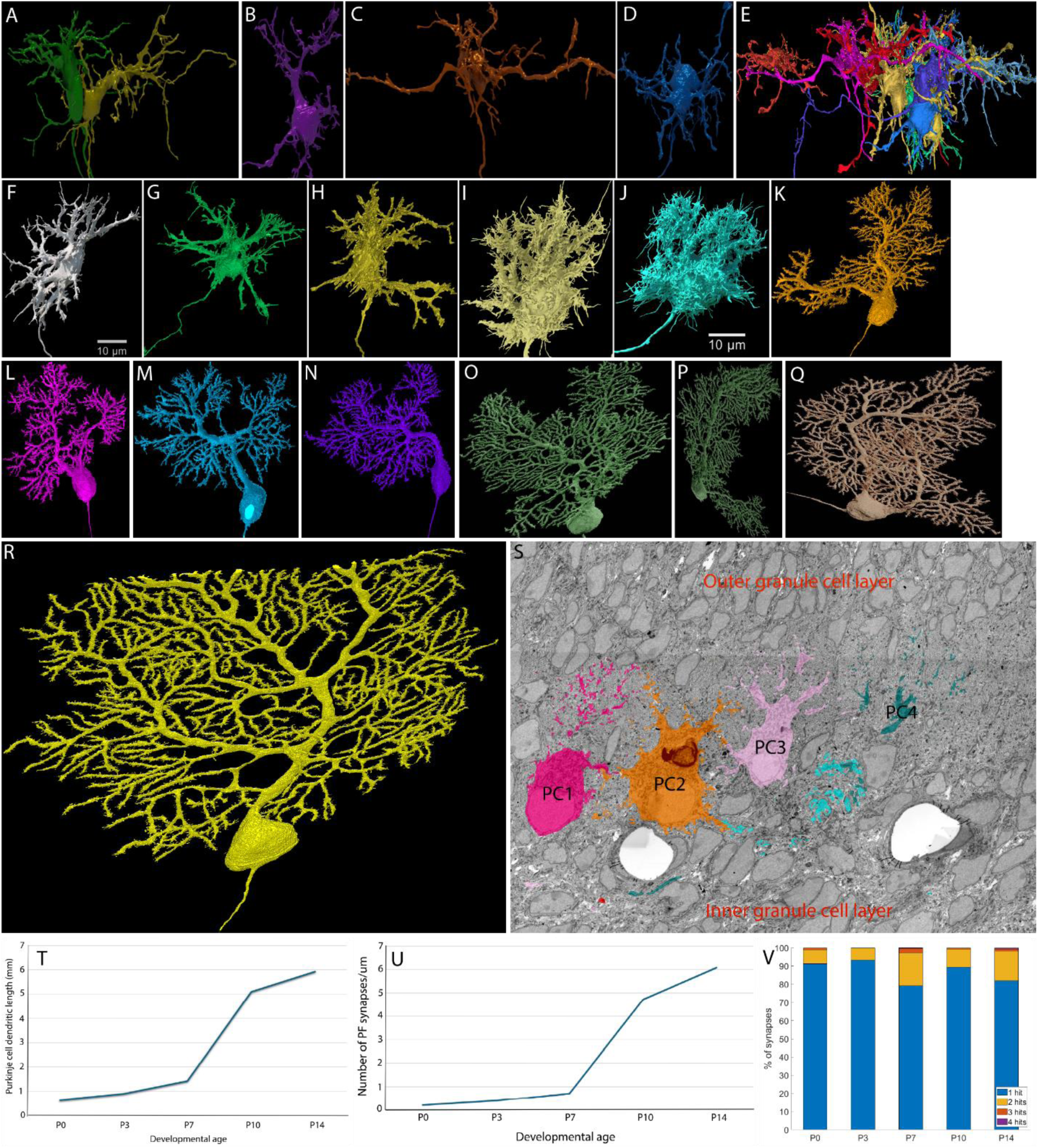
Morphological variability of Purkinje cells within and across developmental ages. Reconstructions of Purkinje cells at P0 A-E, P3 F-H, P7 I-J, P10 K-N, P14 Q-R. Panel E shows a group of P0 Purkinje cells with incoming synapses (yellow circles) on their dendrites. S) Purkinje cells colored (magenta, pink, orange, cyan, and teal) in an electron micrograph of a P7 cerebellar cortex. T) A graph showing the increasing dendritic length of Purkinje cells between P0 and P14. U) A graph showing synaptic density on the Purkinje cell dendrites between P0 and P14. V) A stacked bar graph showing the number of synapses established by a parallel fiber per Purkinje cell during the first two postnatal weeks.

**Figure S3.**
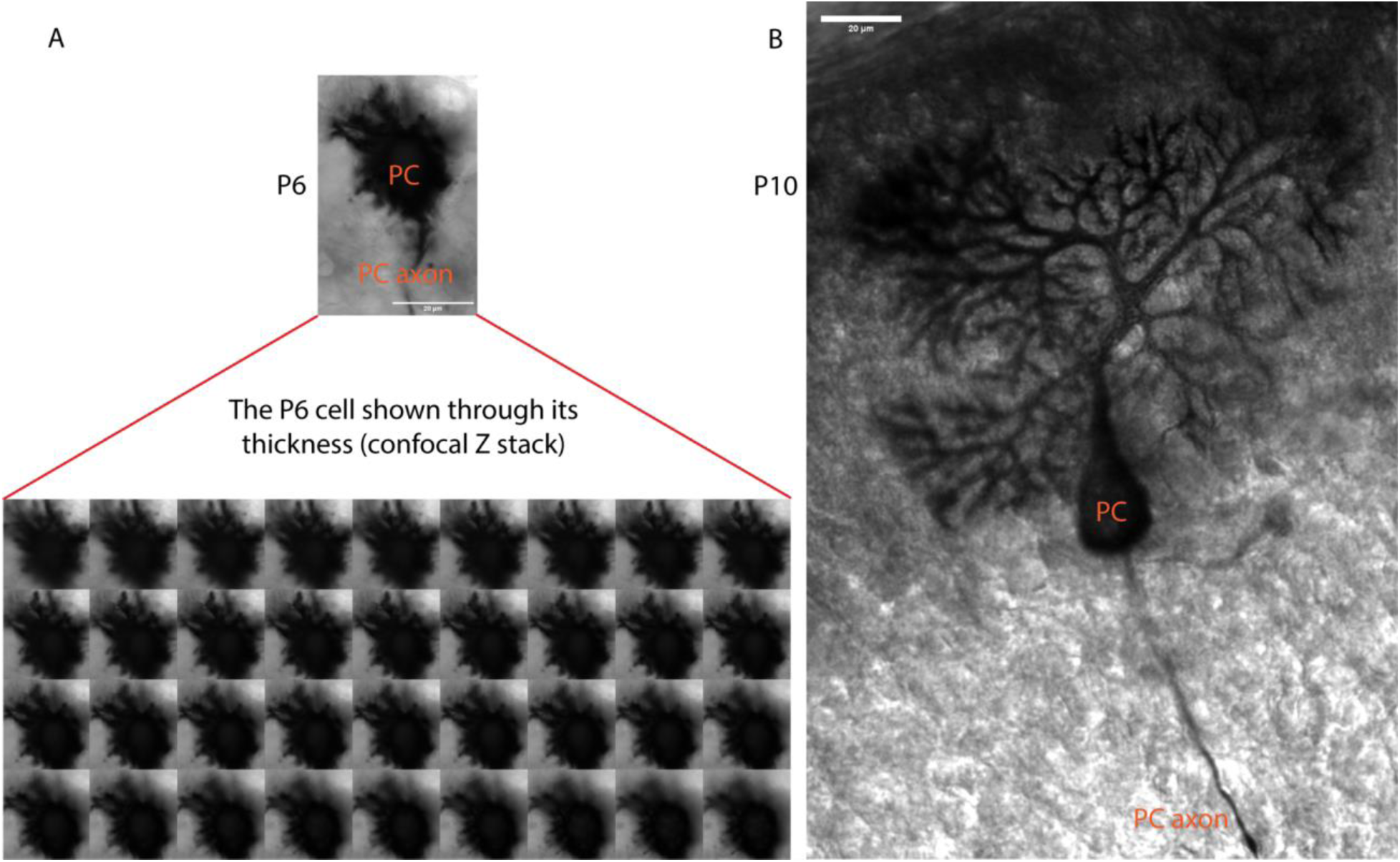
A) A minimum intensity projection of a P6 Purkinje cell imaged using Zeiss LSM 810 Confocal microscope using 40X objective lens (zoom 1.5). PC indicates the cell body of the Purkinje cell and PC axon indicates its axon. All the images of the P6 Purkinje cell at many depths of thickness are shown in a montage below the cell. B) A single image of a P10 Purkinje cell imaged Zeiss LSM 810 confocal using 20X objective. Scale bars in A and B: 20um.

**Figure S4.**
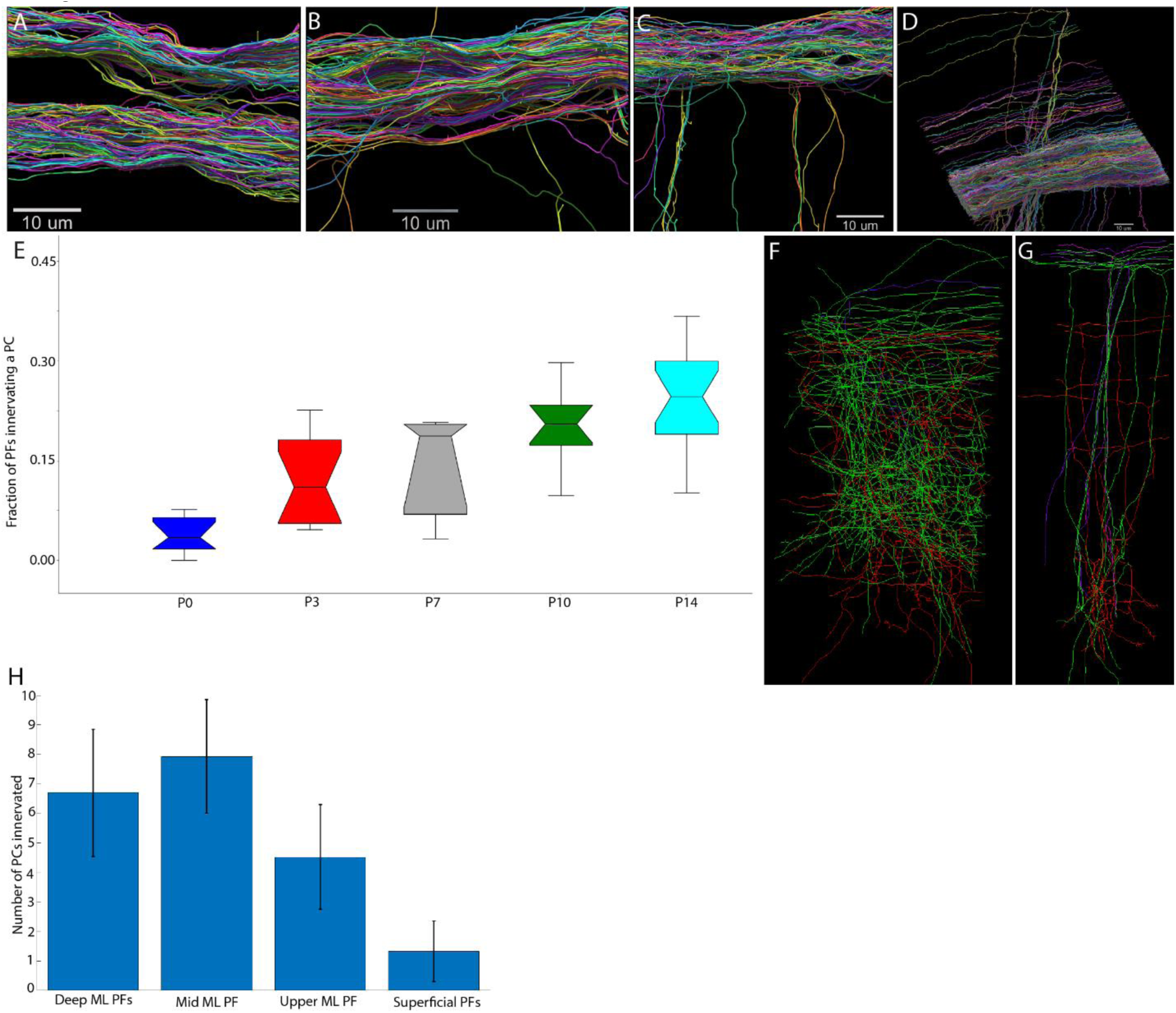
A-D) Reconstructions of dense clusters of parallel fibers at P0 (400), P3 (345), P10 (343) and P14 (2599 axons). E) Skeletonized versions of granule cells showing three distinct connectivity classes. Those that have neither mossy fiber input on their dendrites nor output on Purkinje cells (purple). Those granule cells that received mossy fiber input but have not established parallel fiber output (green). Those that have both input and output (red) at P3, E, and P10 F.

**Figure S5.**
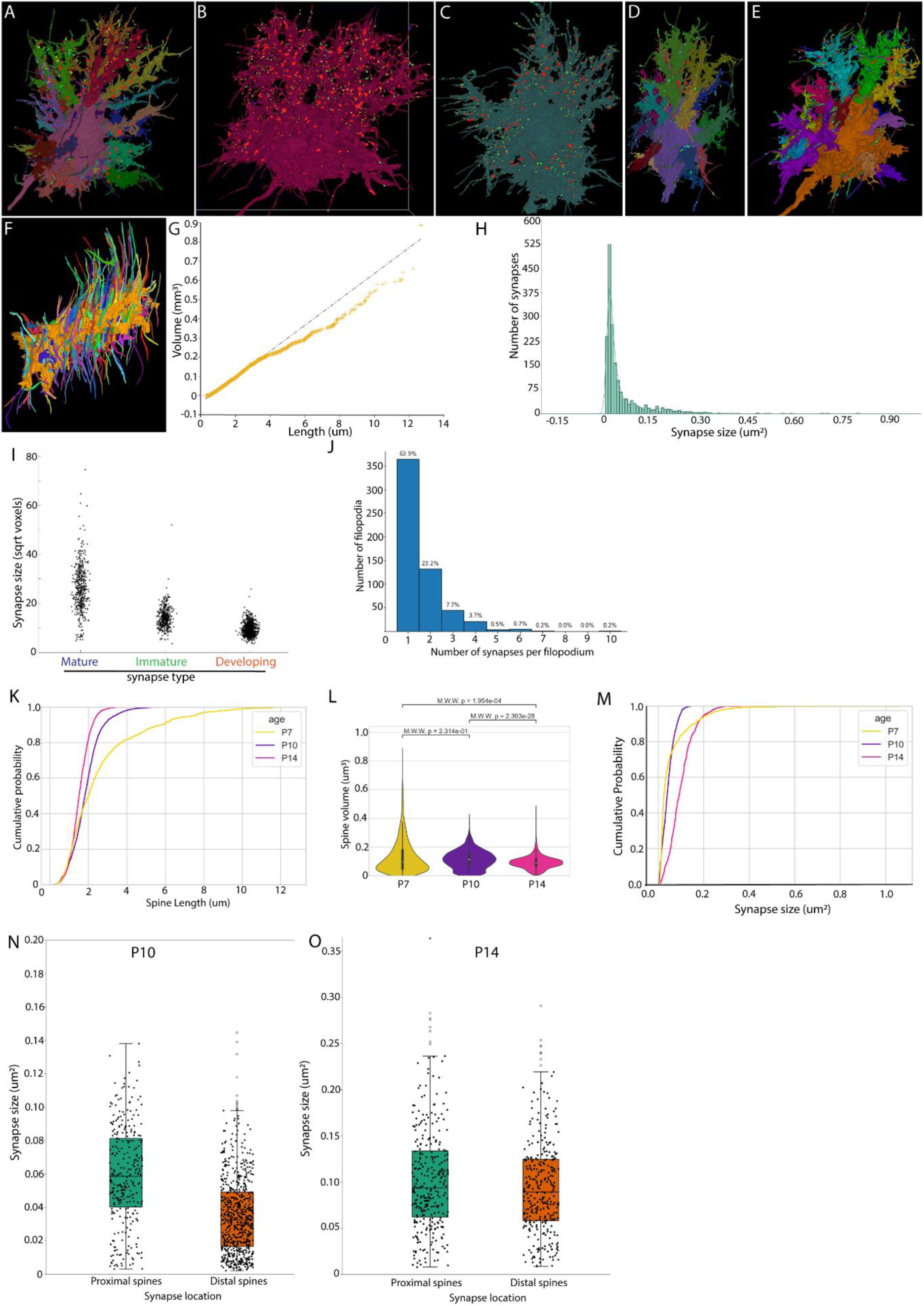
A-e) Shown is the entire complement of all parallel fiber synapses located on filopodia that were labeled for 5 different P7 Purkinje cells, including developing (cyan), immature (green) and mature (red) synapses. F) Filopodia on a dendrite of a P7 Purkinje cell were reconstructed in random colors showing the highly oriented nature of filopodia. G) a graph showing length versus volume of filopodia of P7 Purkinje cells. H) Histogram showing the distribution of synapse sizes present on filopodia with Kernel Density Estimate overlaid. I) Synapses shown in panel H are classified based on the number of vesicles, synaptic contact area and the presence or absence of postsynaptic density. J) A histogram showing number synapses per filopodium on a P7 Purkinje cell. K) A CDF plot shows the lengths of dendritic filopodia at P7 (yellow trace), P10 (purple trace), and P14 (pink trace). L) A combined violin and boxplot showing changes in volume of filopodia at P7 (yellow), P10 (purple) and P14 (pink). M) A CDF plot showing synapse sizes at P7 (yellow trace), P10 (purple trace), and at P14 (pink trace). N-O) A combined box and swarm plot showing comparison of synapse sizes on the spines of proximal (green) versus distal (orange) dendrites of P10 (N) and P14 (O).

**Figure S6.**
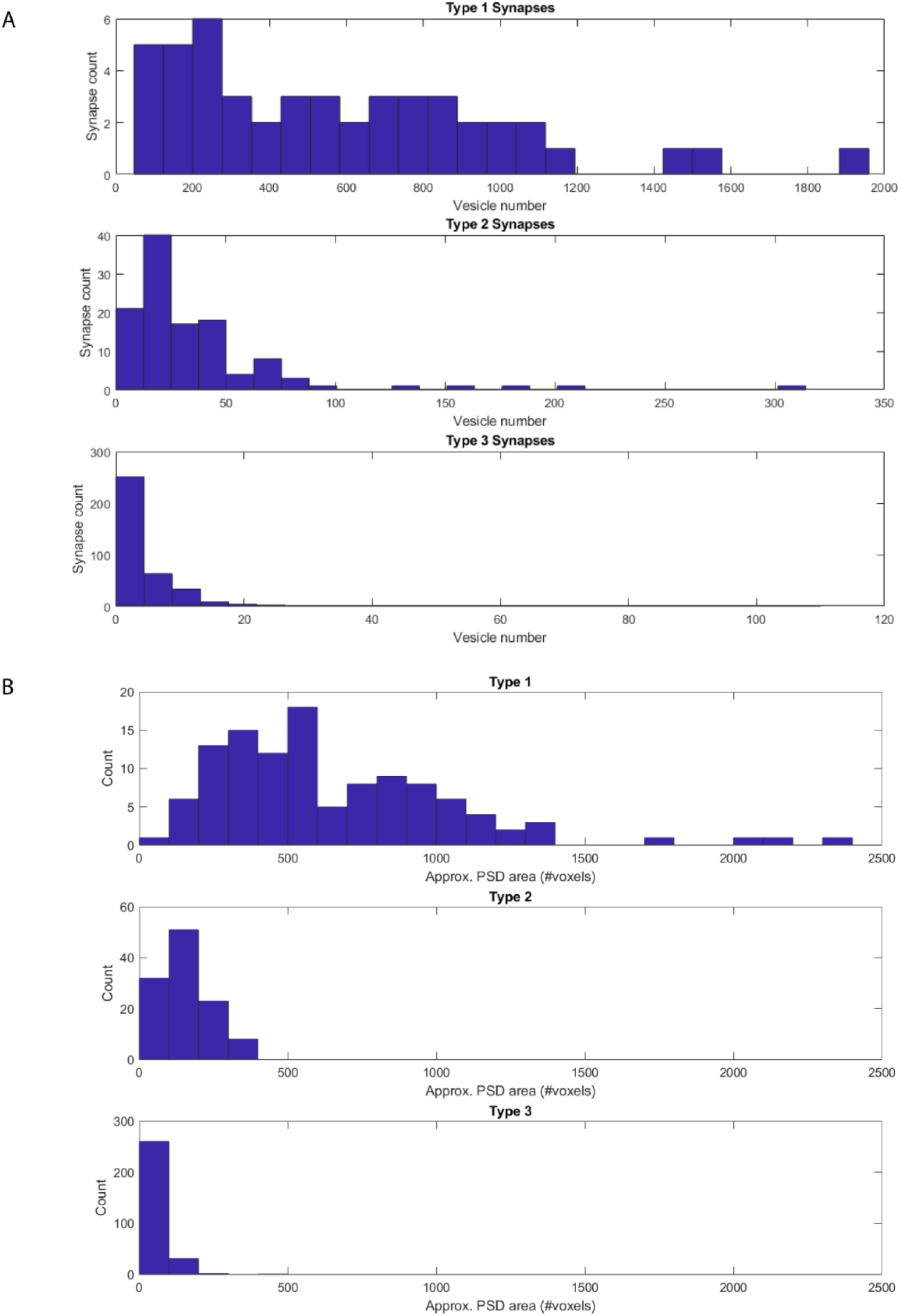
Characterization of parallel fiber synapse maturation at Purkinje cell filopodia. A) A histogram showing the number of synaptic vesicles per filopodial synapse. B) A histogram showing the quantification of area of postsynaptic density measured as voxels labeled.

**Figure S7.**
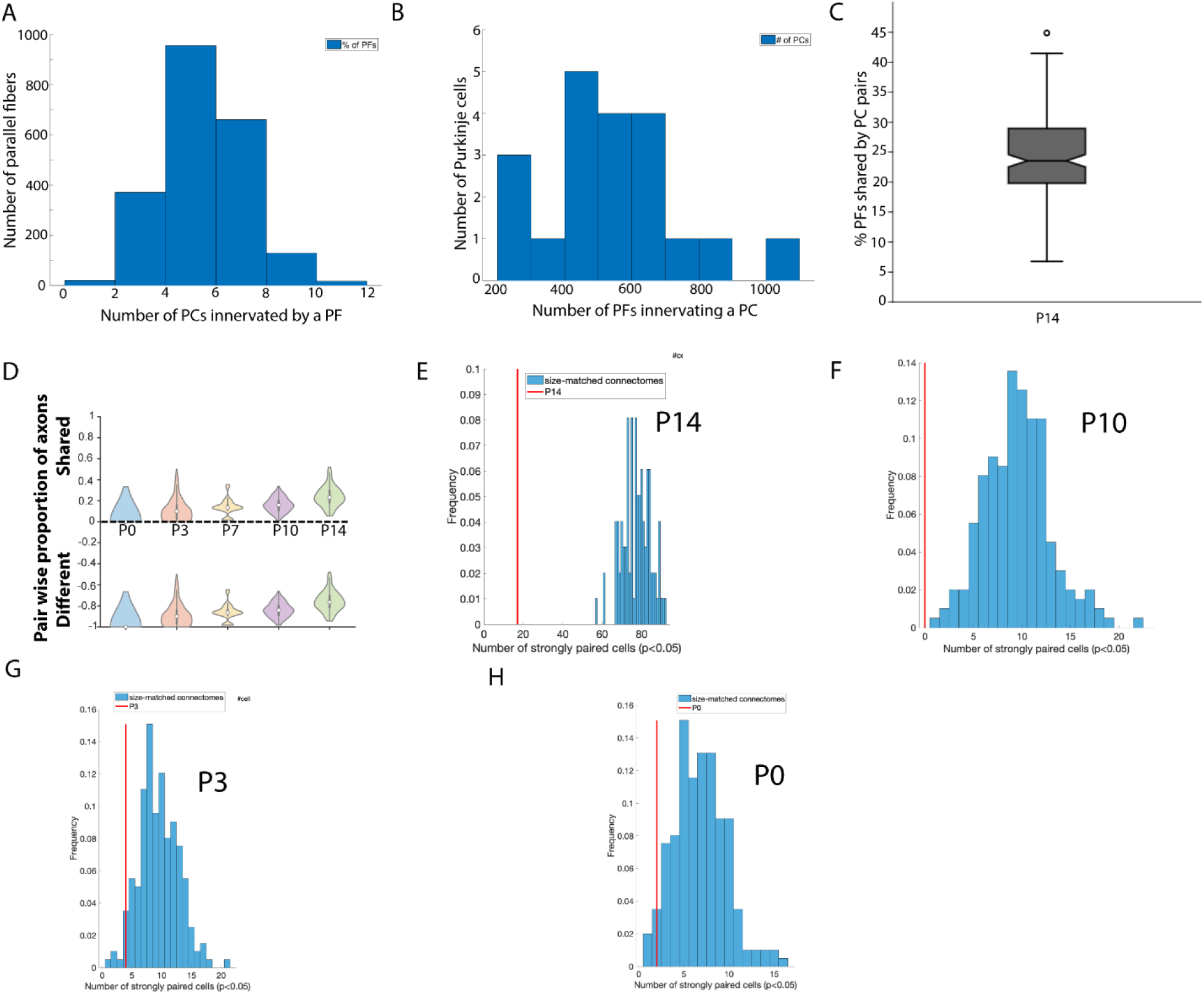
A) Histogram showing the number of Purkinje cells innervated by parallel fibers at P14. B) Histogram of number of parallel fibers innervating a Purkinje cell at p14. C) A box plot showing the percentage of parallel fibers shared by any pair of Purkinje cells (out of 20 PCs analyzed) at P14 (mean, 25%). D) Fraction of parallel fibers shared and not shared (different) by any pair of Purkinje cells at P0, P3, P7, P10 and P14. E-H) Histograms showing comparison of observed connectivity (red line) versus age-matched randomized connectomes (under the constraints of observed connectivity) at P0, P3, P10 and P14.

**Figure S8.**
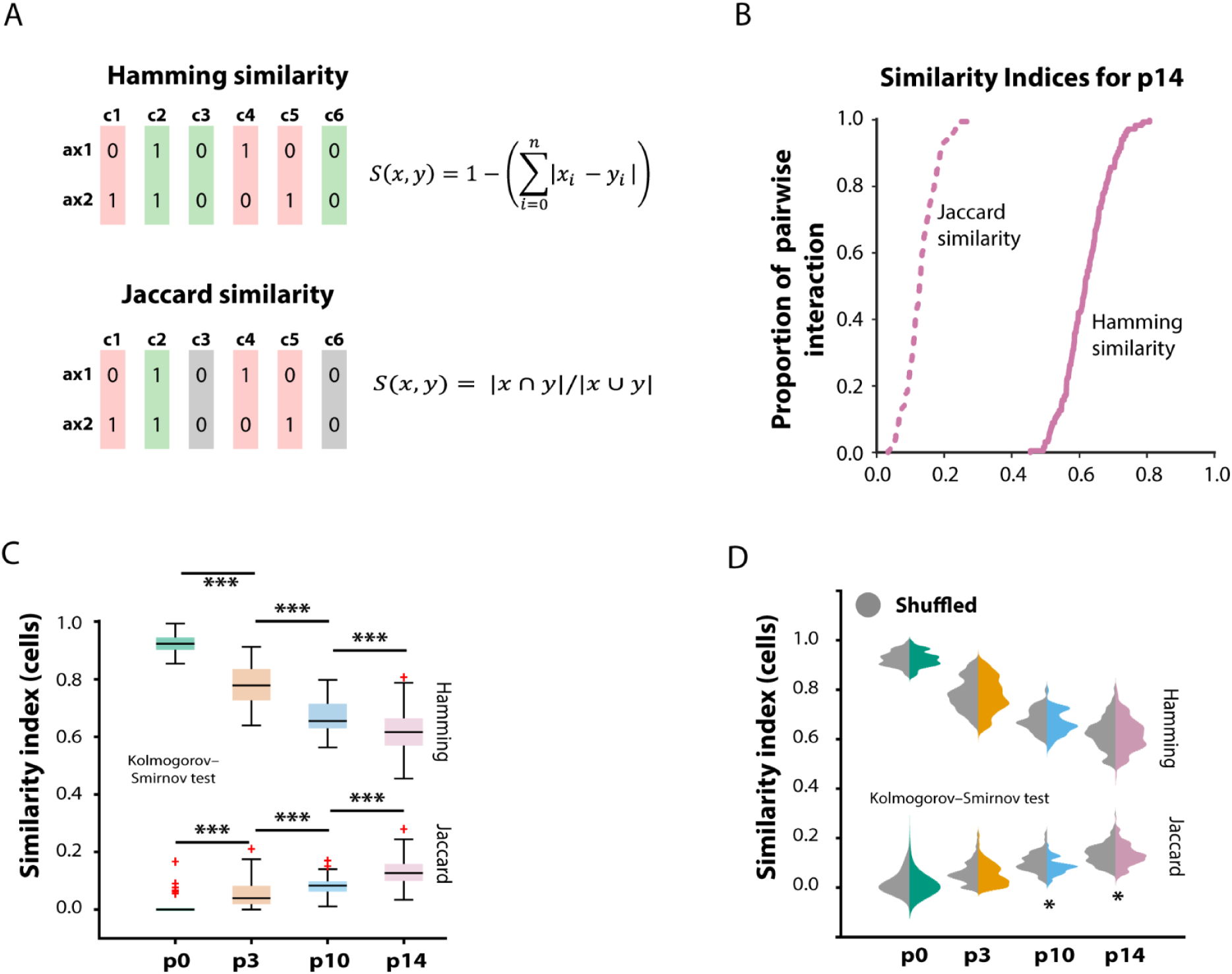
A) Schematic representation and formula of measuring shared parallel fiber connectivity between Purkinje cells using Jaccard and Hamming distances. B) CDF plot showing Jaccard and hamming similarity indices for P14 Purkinje cells. C) Quantification of proportion of shared axons between pairs of Purkinje cells from P0-P14. D) Kullback-Leibler divergence test (left) and Kolmogorov-Smirnov test (right) for both Hamming and Jaccard similarities.

**Figure S9.**
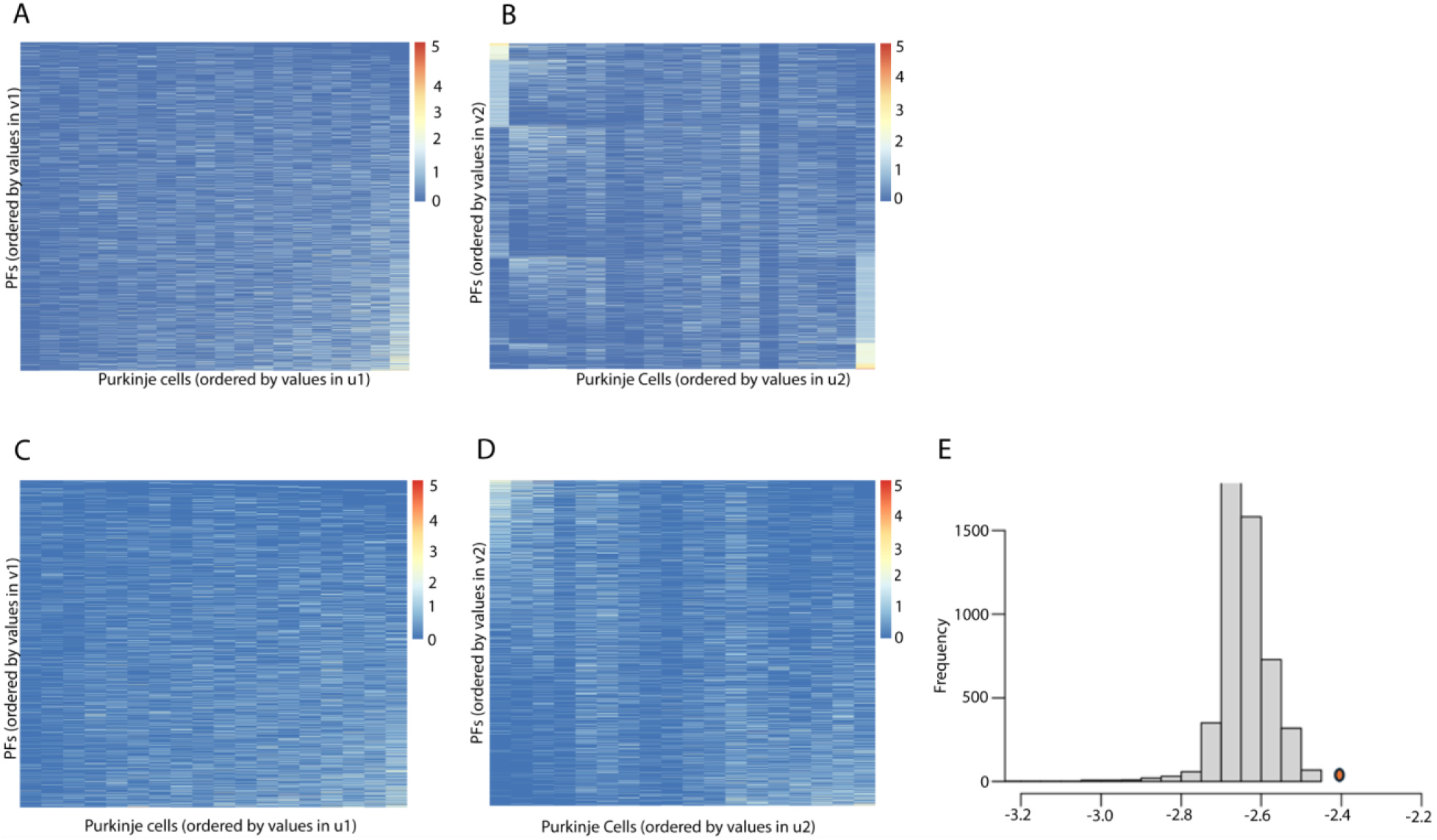
A-B) Heatmaps of P14, where the Purkinje cell and parallel fibers are rearranged based on the values of the latent factors u_i and their cell loadings v_i. In (A), Purkinje cells are roughly based on the total amount of parallel fibers per Purkinje cell, whereas in (B), Purkinje cells and fibers are ordered to highlight the distinct groups of parallel fibers associated with different (groups of) Purkinje cells. C-D) Heatmaps of P14, where the two Purkinje cells corresponding to the first and last columns of the heatmap in (B) are removed, and the remaining cells and parallel fibers are rearranged based on u_i and v_i recomputed based on the remaining cells. In (C), Purkinje cells are roughly based on the total amount of parallel fibers per Purkinje cell, whereas in (D), Purkinje cells and fibers are ordered to highlight that the remaining Purkinje cells are more evenly associated with the parallel fibers, that is, different cells tend to connect with distinct fibers rather than share the same group of fibers. E) Histogram of the simulated null distribution of the entropy measure for the second latent factor. The red dot marks the observed entropy value for the second latent factor of P14 after removing the two Purkinje cells. A significantly elevated entropy relative to the null indicates strong evidence for an even distribution of fibers across cells that is unlikely to have occurred by chance.

